# Orderly assembly underpinning built-in asymmetry in the yeast centrosome duplication cycle requires cyclin-dependent kinase

**DOI:** 10.1101/2020.05.22.110296

**Authors:** Marco Geymonat, Qiuran Peng, Zhiang Guo, Zulin Yu, Jay R. Unruh, Sue L. Jaspersen, Marisa Segal

**Affiliations:** Stowers Institute for Medical Research, Kansas City, MO 64110; Department of Molecular and Integrative Physiology, University of Kansas Medical Center, Kansas City, KS 66160, USA; Department of Genetics, University of Cambridge, Downing Street, Cambridge CB2 3EH, UK

**Keywords:** spindle, polarity, asymmetric fate, *S. cerevisiae*, gamma-tubulin complex

## Abstract

Asymmetric astral microtubule organization drives the polarized orientation of the *S. cerevisiae* mitotic spindle and primes the invariant inheritance of the old spindle pole body (SPB, the yeast centrosome) by the bud. This model has anticipated analogous centrosome asymmetries featuring in self-renewing stem cell divisions. We previously implicated Spc72, the cytoplasmic receptor for the gamma-tubulin nucleation complex, as the most upstream determinant linking SPB age, functional asymmetry and fate. Here we used structured illumination microscopy and biochemical analysis to explore the asymmetric landscape of nucleation sites inherently built into the spindle pathway and under the control of cyclin-dependent kinase (CDK). We show that CDK enforces Spc72 asymmetric docking by phosphorylating Nud1/centriolin. Furthermore, CDK-imposed order in the construction of the new SPB promotes the correct balance of nucleation sites between the nuclear and cytoplasmic faces of the SPB. Together these contributions by CDK inherently link correct SPB morphogenesis, age and fate.

## INTRODUCTION

Spindle orientation in self-renewing stem cell divisions exploits structural asymmetries built into the centrosome cycle to create a directional bias that links differential fate with an invariant pattern of age-dependent centrosome inheritance (Fu et al., 2015; Pelletier and Yamashita, 2012; Rebollo et al., 2007; Venkei and Yamashita, 2018; Wang et al., 2009; Yamashita et al., 2007). Perturbation of these mechanistic links impairs self-renewal, prompting an imbalance between stem cell pools and differentiating progeny that disrupts development or causes tumourigenesis (Gonzalez, 2013; Mukherjee and Brat, 2017; Vertii et al., 2018; Wang et al., 2009; Wodarz and Nathke, 2007).

The premise of an invariant pattern of spindle pole inheritance coupled to spindle orientation in cells dividing asymmetrically first emerged in the budding yeast *Saccharomyces cerevisiae* (Pereira et al., 2001), a unicellular organism that divides into a larger mother cell and a smaller daughter cell or bud. In *S. cerevisiae*, all aspects of spindle morphogenesis are controlled by the spindle pole body (SPB), the analog of the animal centrosome (Byers, 1981; Cavanaugh and Jaspersen, 2017; Fu et al., 2015; Jaspersen and Winey, 2004; Winey and Bloom, 2012). The SPB consists of three major layers — an inner plaque facing the nucleus, a central plaque rooted in the nuclear envelope and an outer plaque facing the cytoplasm (Figure 1 A and Supplementary Figure 1A). A specialized extension on one side, the half-bridge, is required for SPB duplication (Byers and Goetsch, 1975). Microtubule (MT) nucleation sites arise by recruitment of the conserved *γ*-tubulin complex (*γ*TC but also referred to as the *γ*-tubulin small complex or *γ*-TuSC) composed of Tub4 (yeast *γ*-tubulin), Spc98/GCP3 and Spc97/GCP2 (Farache et al., 2018; Geissler et al., 1996; Knop et al., 1997; Kollman et al., 2011; Winey and Bloom, 2012). The inner plaque organizes intranuclear spindle MTs by docking *γ*TCs onto Spc110/pericentrin (Knop and Schiebel, 1997). By contrast, nucleation of cytoplasmic or astral microtubules (aMTs) occurs from two distinct *γ*TC-docking sites set up by the cytoplasmic receptor Spc72/CDK5RAP2, upon binding Nud1/centriolin at the outer plaque or Kar1 at the half-bridge (Gruneberg et al., 2000; Knop and Schiebel, 1998; Lin et al., 2015; Pereira et al., 1999).

**Figure 1.**
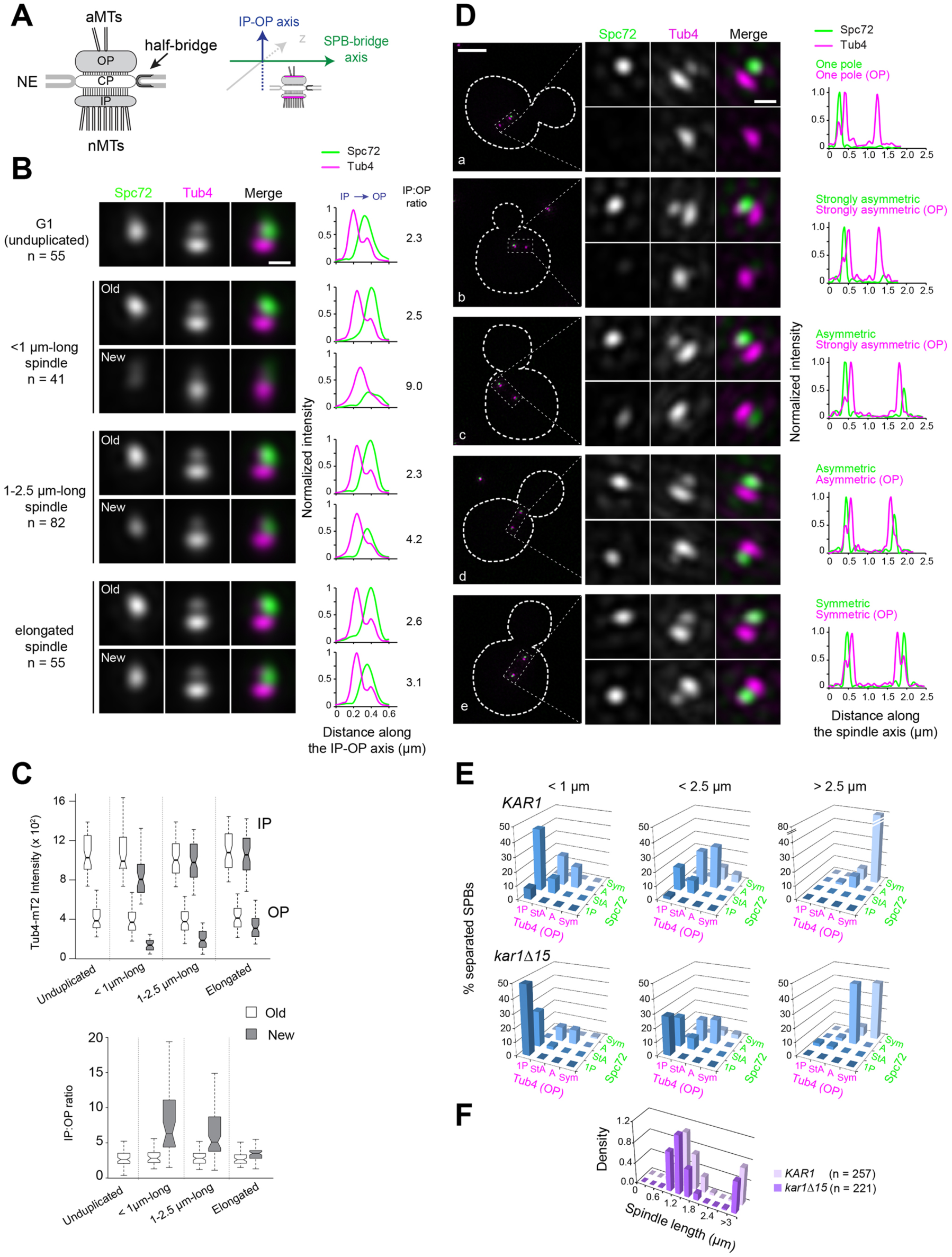
Tub4 accumulation at SPB outer plaques follows Spc72 asymmetric recruitment during spindle assembly. (A) [Left] Simplified schematic of the SPB showing the inner (IP) and outer (OP) plaques that nucleate nuclear (nMTs) and cytoplasmic or astral (aMTs) microtubules, respectively. The central plaque (CP) anchors the SPB in the nuclear envelope (NE). The half-bridge is a modified region of the nuclear envelope involved in SPB duplication; it is also able to nucleate microtubules during G_1_ phase of the cell cycle (Byers & Goetsch 1975). [Right] To study the distribution of SPB components, SIM images of individual SPBs were aligned using Tub4-mT2 as a reference, as previous work showed that it is present at the inner and outer plaque in different amounts (see Burns et al., 2015 and Supplementary Figure 1). SPBs were assigned to a cell cycle and spindle stage using bud morphology and the distances between SPBs, referred to as SPB inter-distances. (B) Average image from realigned SPBs in each class. Scale bar, 200 nm. The number of SPBs is indicated. Linescan analysis (5-px width) shows the intensity and localization of Spc72-Venus as well as the distribution of Tub4-mT2 at the IP and OP. Linescan intensities were normalized relative to maximal values at the old SPB of elongated spindles. The IP:OP ratio is based on the Tub4-mT2 signal. (C) In addition to averaging, the intensity of Tub4-mT2 at the inner and outer plaques was measured in individual images. Values are plotted by stage and SPB identity (top); IP:OP intensity ratios (bottom) for the same dataset are also shown. Boxplots depict the 5th, 25th, 50th, 75th and 95th centiles. Notches represent 95% CI of the median. (D) Representative SIM images showing the distribution of Spc72-Venus (green) and Tub4-mT2 (magenta) at SPBs of cells with short spindles. Merged images with cell outlines (scale bar, 2 µm) and single channels and merged cropped images for each SPB (scale bar, 200 nm) are shown paired with internally normalized linescan analysis to indicate fluorescence intensity along the spindle axis: (a) Spc72 and Tub4 present at one outer plaque; (b-d) progressive decline of asymmetry for Spc72 followed by Tub4; (e) both components symmetric. (E) Quantitation of Spc72-Venus and Tub4-mT2 distribution by spindle stage in *KAR1* and *kar1Δ15* strains. 1P = one pole; StA = strongly asymmetric; A = asymmetric; Sym = symmetric, as described in D. (F) Distribution of spindle lengths in the *KAR1* and *kar1Δ15* asynchronous cell populations analyzed.

Conservative SPB duplication generates a new SPB next to the old SPB inherited from the previous cell cycle, thus laying the foundations for inherent functional asymmetry linked to SPB age. Genetic analyses in combination with electron microscopy (EM) and, more recently, structured illumination microscopy (SIM), have contributed toward a model mapping the ordered addition of individual components according to three landmark events — satellite assembly at the distal end of the bridge, generation of a duplication plaque and a final side-by-side stage with the bridge connecting duplicated SPBs (Adams and Kilmartin, 1999; Burns et al., 2015; Cavanaugh and Jaspersen, 2017; Jaspersen and Ghosh, 2012; Jaspersen and Winey, 2004; Ruthnick and Schiebel, 2018; Winey and Bloom, 2012). By onset of SPB separation and spindle assembly, the old SPB has established first contacts with the bud through existing aMTs while the newly assembled SPB delays aMT organization until a ∼1 µm-long spindle has formed (Shaw et al., 1997). This intrinsic functional asymmetry, in interplay with extrinsic cues, primes spindle polarity and orientation along the mother-bud axis in association with the stereotyped inheritance of the old SPB by the bud (Geymonat and Segal, 2017). In agreement with live imaging data (Juanes et al., 2013; Segal et al., 2000; Shaw et al., 1997), three-dimensional ultrastructural analyses demonstrate aMT asymmetric organization built into the early stages in the spindle pathway, with aMTs emerging from the old SPB outer plaque and the bridge both at the satellite and side-by-side stages in cells proceeding unperturbed (Byers and Goetsch, 1975; McIntosh and O’Toole, 1999; O’Toole et al., 1999).

Core cell cycle controls linking aMT organization with landmark events along the spindle pathway might involve phosphorylation targets at the SPB. Phosphorylation sites have been identified in many SPB components but their possible significance to intrinsic SPB functional asymmetry remains unknown (Fong et al., 2018; Huisman et al., 2007; Keck et al., 2011; Lin et al., 2011; Lin et al., 2014; Rock et al., 2013). In *S. cerevisiae*, cell cycle progression is controlled by a single cyclin-dependent kinase (CDK), Cdc28/Cdk1, that associates with a series of phase-specific cyclins to control various cell cycle events (Morgan, 2007). We have previously implicated the S-phase CDK Clb5-Cdc28 in enforcing aMT asymmetry. Indeed, Clb5 inactivation in the sensitized *cdc28-4* background, *cdc28-4 clb5Δ,* specifically abrogates the delay in aMT organization at the new SPB relative to spindle assembly with concomitant disruption of spindle polarity (Segal et al., 2000; Segal et al., 1998). More recently we have correlated aMT temporal asymmetry with the acquisition of Spc72 at the new SPB outer plaque during spindle assembly (Juanes et al., 2013). However, that study could not determine how Spc72 temporal asymmetry arises or its direct impact on *γ*TC distribution at distinct nucleation sites. Moreover, the idea that intrinsic SPB asymmetry has such structural basis has been brought into question in a recent study (Lengefeld et al., 2018).

Here we undertook detailed analysis of the mode of distribution of Spc72 and components of the *γ*TC throughout the spindle pathway by SIM. Our data show that biased recruitment of Spc72 and the *γ*TC at the old SPB stems from manifest asymmetry throughout G_1_ and during the side-by-side stage, which is preserved until spindle assembly begins. Our data also show that CDK targets Nud1 in order to enforce Spc72 biased partition during S phase. Moreover, CDK is required for the *γ*TC inner to outer plaque ratio that underlies the normal distribution of nucleation sites at the two faces of the SPB. Finally, CDK also promotes the assembly of the new SPB in the correct order by which *γ*TC addition to the outer plaque always follows the complete assembly of the inner plaque. Taken together, our study reveals key contributions of CDK toward accurate SPB morphogenesis that secure the correct interplay between intrinsic and extrinsic asymmetries that determine spindle polarity and differential fate.

## RESULTS

### Asymmetric outer plaque structure and γTC partition during spindle assembly revealed by SIM

We have previously proposed that the intrinsic link between SPB age, temporally asymmetric aMT organization and SPB fate is underpinned by Spc72 biased localization to the old SPB during spindle assembly (Juanes et al., 2013). Yet, how this asymmetry is built and the associated impact on the localization of *γ*TC at three nucleation sites — inner plaque, outer plaque and the bridge — remain unknown. SIM represents an effective approach to answer these questions as demonstrated by a previous study focused on core molecular events in SPB duplication (Burns et al., 2015). Of particular relevance is the ability of SIM to resolve SPB inner and outer plaques shown by that study, using strains expressing endogenous SPB components fused to fluorescent tags. Accordingly, single SPBs of cells expressing Tub4-mTurquoise2 (mT2) showed two distinct foci that could be assigned to inner and outer plaques by co-label with YFP-Spc110 or YFP-Spc72, respectively (Burns et al., 2015 and Supplementary Figure 1 B). Two foci were also observed using Spc97-mT2 or Spc98-mT2 (Burns et al. 2015). In all cases, the focus co-localized with YFP-Spc110 was more intense than the signal co-localized with YFP-Spc72, consistent with the idea that there is more *γ*TC at the inner plaque of the SPB than at the outer plaque.

To establish the relative ratio of *γ*TC at the inner and outer plaque, we determined the intensity of Tub4-mT2, Spc97-mT2 or Spc98-mT2 foci in individual SIM images (Supplementary Figure 1 C). Based on the accepted nuclear to cytoplasmic MT ratio of ∼6-7 (Erlemann et al., 2012), we anticipated a similar ratio for the *γ*TC. Against expectations, quantitative analysis uncovered an overall inner to outer plaque ratio (IP:OP ratio) of ∼ 2-3 for each *γ*TC component. While it is clear from individual measurements there is considerable heterogeneity in *γ*TC that will be further investigated below, this heterogeneity, the lower z-resolution of SIM or other issues with quantitation are unlikely to account for the observed IP:OP ratio. In diploid cells, the number of MTs increases two-fold (Byers and Goetsch, 1974). Analysis of Tub4-mT2 distribution in haploids and diploids showed a roughly two-fold increase in *γ*TC at both the inner and outer plaques of diploid SPBs (Supplementary Figure 1 D). Collectively, these data point to an unexpected, marked contribution of the outer plaque to bulk *γ*TC content at the SPB and illustrated the utility of SIM for analysis of SPB structure. This finding is consistent with the notion that Tub4 asymmetry observed by wide-field fluorescence microscopy might reflect events at the cytoplasmic face of the SPB (Juanes et al., 2013).

The *γ*TC localizes to two cytoplasmic SPB substructures: the outer plaque and the (half)-bridge. To better understand possible *γ*TC heterogeneity and *γ*TC distribution to the outer plaque or half-bridge as contributors to intrinsic SPB asymmetry, we analyzed Tub4-mT2 by SIM in asynchronous cell populations in either *KAR1* or *kar1Δ15* backgrounds. The *kar1Δ15* mutation abrogates Spc72 binding to Kar1 (Pereira et al., 1999), thus precluding *γ*TC localization to the bridge. Strains also contained Spc72-Venus so that images in which inner and outer plaques overlapped vertically could be excluded (top view), an issue with previous single color SIM analysis of *γ*TC (Lengefeld et al., 2018). Judging from the apparent overlap between single Tub4 foci and the core outer plaque component Nud1, only ∼ 12% of SPBs in our SIM images showed a top view (Supplementary Figure 1 E).

Cell images were grouped by cell cycle/spindle stage (SPB number and spindle length) and SPB age (old versus new) based on measure of inner plaque intensities and/or position within the cell. To compare Tub4 localization among multiple SPBs in a cell cycle/spindle stage, we used computational methods to align dual color SIM images based on Tub4-mT2 fluorescence at the inner and outer plaque of SPBs within that category and used to generate an averaged image (Figure 1 A-B, see Materials and Methods). This type of analysis is advantageous because it shows the likelihood that a protein is present in a given location based on many cells and allows for positional comparison between different proteins (Burns et al., 2015). From these averaged images and their quantitation by linescan analysis (Figure 1 B) or from quantitation of individual SPBs (Figure 1 C), the structural asymmetry between old versus new SPB was apparent. Unduplicated SPBs from G_1_-phase cells exhibited a Tub4 IP:OP ratio between 2 and 3 that was also manifest in old SPBs throughout the remaining of the cell cycle. By contrast, the ratio was highest in new SPBs early in spindle assembly but decreased later in the spindle pathway.

Linescan analysis along the spindle axis in maximal SIM image projections of wild type and *kar1Δ15* cells was used to elucidate the contribution of the bridge pool of γTC to its asymmetry (Figure 1 D-F), validating our data from averaged images while revealing further significant trends. First, Tub4-mT2 was apparent at the inner plaque of the new SPB early in spindle assembly (< 1 µm-long spindles), before Spc72-Venus or Tub4-mT2 accumulated at the new SPB outer plaque (Figure 1 D a-b). Both components were strongly asymmetric in *KAR1* cells or mainly present at the old pole in *kar1Δ15* cells (Figure 1 E). The complete assembly of the SPB inner plaque served as a further reference in support of the late acquisition of Tub4 by the new SPB outer plaque. Second, incorporation of Tub4-mT2 onto the new SPB trailed Spc72-Venus, consistent with the idea that Spc72 is needed for γTC recruitment to the outer plaque (Figure 1 E). Taken together, Tub4 exhibed a marked asymmetry at the cytoplasmic face of the SPB at early stages of the spindle pathway underpinned by Spc72 distribution. Further, the bridge-based localization of γTC plays little role in this asymmetry.

Importantly, Spc72 and Tub4 asymmetry during spindle assembly was also observed in the W303 yeast strain background in both individual images and in averaged images grouped by cell cycle/spindle stage (Supplementary Figure 2 A-C). Quantitative analysis for relative Tub4 partition between inner and outer plaques by stage and SPB identity confirmed that Tub4 asymmetric recruitment is common to both 15D and W303 strain backgrounds (Supplementary Figure 2 B-C). Validation of Spc72 asymmetry by wide-field fluorescence microscopy analysis in multiple strains (Supplementary Figure 2 D) indicates that the effects are unlikely related to yeast genetic background and/or SIM. Rather, these results demonstrate that intrinsic SPB structural and functional asymmetry is a general feature of the budding yeast spindle cycle.

### Spc72 and Tub4 asymmetry arise during SPB duplication

Landmark molecular events in SPB duplication have been elucidated by EM studies and, more recently, dissected by SIM (Adams and Kilmartin, 1999; Burns et al., 2015). In order to understand the source of SPB functional asymmetry at onset of spindle assembly, we focused on characterizing events at the cytoplasmic face of the SPB(s) along the duplication pathway, with emphasis on the distribution and incorporation of Spc72 and γTC, not studied in previous SIM analysis. However, it would be tempting to predict the temporality for assembly of the outer plaque on the basis of prevalent models for SPB duplication (Figure 2 A; Winey and Bloom, 2012).

**Figure 2.**
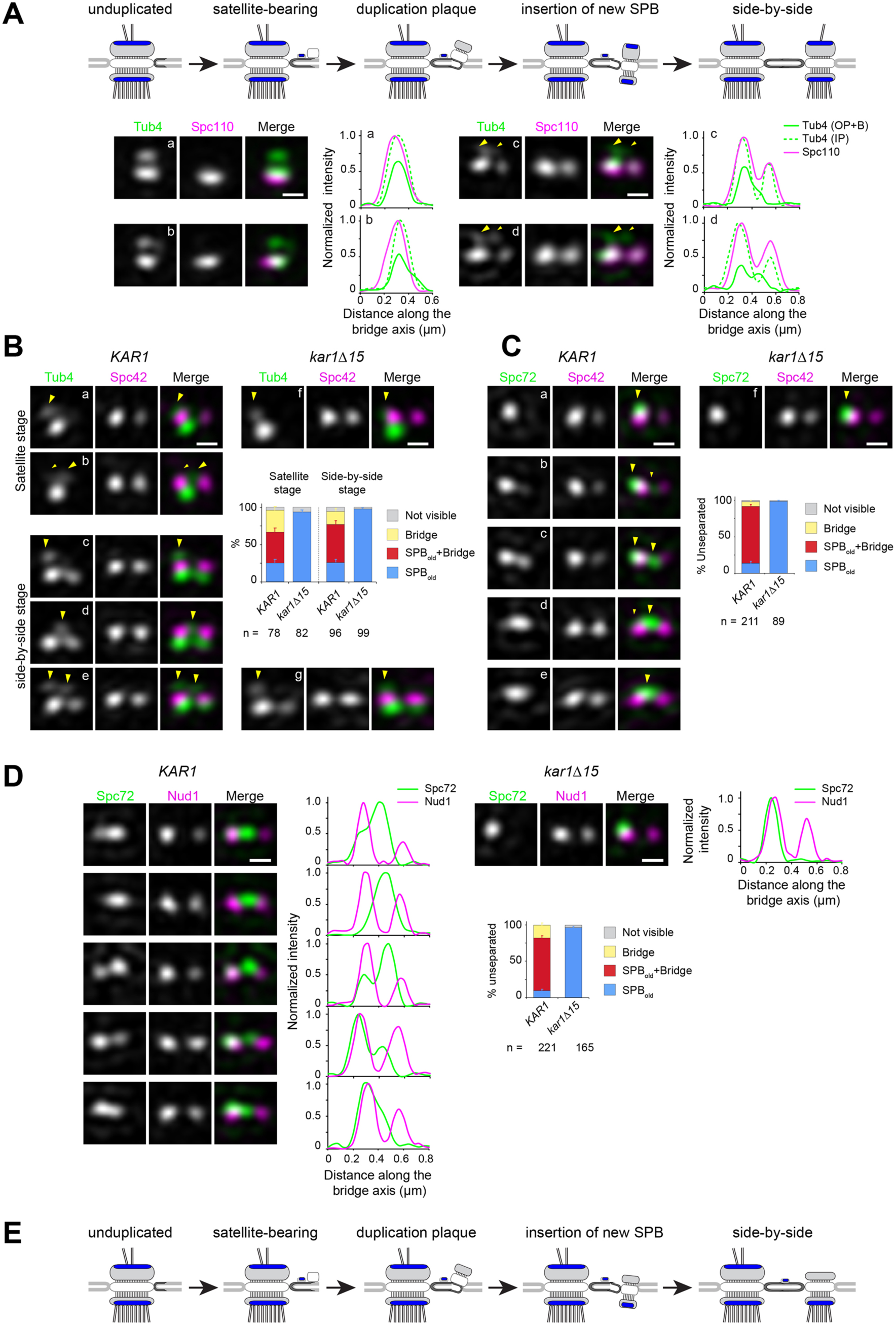
Spc72 and γTC are not added to the new SPB during its duplication. (A-D) SIM image projections were obtained after 3D-Gaussian fit and realignment using the indicated reference label. Images were rotated to position the old SPB to the left. Scale bar, 200 nm. (A) (top) Model for landmark events along the SPB duplication pathway deduced by EM and SIM studies (after Adams and Kilmartin, 1999 and Burns et al., 2015). The presumed localization of *γ*TC according to this model is suggested in blue. (bottom) Representative SIM images and corresponding linescan analysis along the inner or outer plaque (1px-width, parallel to the bridge axis, internally normalized) illustrating Tub4-Venus distribution (green) relative to mT2-Spc110 (magenta). Images are ordered along the presumed SPB duplication pathway according to structures seen in early or intermediate steps of SPB duplication leading to the completed side-by-side SPBs: single SPB showing either a single (a) or extended (b) cytoplasmic label; (c-d) side-by-side stage. (B-C) Using Spc42-mT2 intensity, SIM images were organized to represent stages of SPB duplication. (B) The distribution of Tub4-Venus (green) localization relative to Spc42-mT2 (magenta) during SPB duplication in *KAR1* (a-e) versus *kar1Δ15* (f-g) cells and plot for distribution of modes of Tub4 localization at the cytoplasmic face of the SPB. (C) The distribution of Spc72-Venus (green) localization relative to Spc42-mT2 (magenta) during SPB duplication in *KAR1* (a-e) versus *kar1Δ15* (f) cells and plot for modes of Spc72-Venus distribution at unseparated SPBs. Error bars, standard error of the proportion. (D) Spc72-Venus (green) and Nud1-mT2 (magenta) were localized in *KAR1* versus *kar1Δ15* cells by SIM. Representative images of cells undergoing SPB duplication, paired to linescan analysis (3 px-width along the bridge axis; internally normalized) and plot for distribution of modes of Spc72 localization relative to Nud1. Error bars, standard error of the proportion. (E) New model depicting the temporality for completion of inner and outer plaque assembly relative to other landmark events in the SPB duplication cycle as emerging from the data presented here. In particular, the new SPB outer plaque is incompletely assembled at the side-by-side stage, and thus unable to recruit *γ*TC.

From asynchronous wild type cells coexpressing mT2-Spc110 (to discriminate inner and outer plaque) and Tub4-Venus, we could identify multiple configurations of γTC in G_1_ cells. In early G_1_, unduplicated SPBs contained two Tub4-Venus foci, the strongest corresponding to the inner plaque (Figure 2 A, a). A subset of unduplicated SPBs exhibited a cytoplasmic signal extending beyond the presumed outer plaque (Figure 2 A, b); this most likely corresponded to bridge-localized γTC. In later G_1,_ two foci of Spc110 were apparent (Figure 2 A, c-d) along with a second prominent Tub4 focus at the new inner plaque, consistent with the idea that Spc110 incorporation occurs after SPB insertion into the nuclear envelope and that Spc110 recruits γTC to the SPB inner plaque. Duplicated SPBs remained unseparated in S phase (see Supplementary Figure 3 A and B for a distribution of distances between unseparated SPB pairs) and had a variable cytoplasmic label of Tub4-mT2 extending from the old SPB to the space between the two inner plaques, as shown in the 1-pixel linescans across inner and outer plaques (Figure 2 A, c-d). These data suggest that γTC is present on the outer plaque of the old SPB throughout SPB duplication and localizes to varying degrees to the bridge. However, it does not appear to be added to the outer plaque of the new SPB as part of SPB duplication in G_1_.

To quantify the distribution of γTC at the old SPB, bridge and new SPB, we compared Tub4-Venus to Spc42-mT2 in wild type or *kar1Δ15* cells. Spc42 is the first protein incorporated into the new SPB during its formation in G_1_ phase, and its levels can be used to evaluate progression through SPB duplication (Adams and Kilmartin, 1999; Burns et al., 2015; Ruthnick et al., 2017). Cells at the satellite or duplication plaque stage were recognized by the presence of duplicated Spc42 signals alongside a single Tub4 inner plaque focus (Figure 2 B, a-b) or at the side-by side stage by the presence of duplicated Spc42 signals with two Tub4 inner plaque foci (Figure 2 B, c-e). During the satellite stage, Tub4 was observed at the old SPB outer plaque (Figure 2 B, a, arrowhead), or favoring the space between the SPB and satellite with a residual signal often apparent at the old SPB outer plaque (Figure 2 B, b, big and small arrowheads, respectively). After appearance of the new inner plaque (Figure 2 B, c-e), label continued to distribute between those two locations. Label associated with the new SPB outer plaque was not observed. In *kar1Δ15* cells Tub4 retained localization exclusively at the old SPB outer plaque throughout (Figure 2 B, f-g) demonstrating that localization at the center of the SPB pair in *KAR1* cells represented nucleation sites at the bridge. Thus, Tub4 and by extension γTC, localizes to the old SPB and bridge during SPB duplication. The fact that Tub4 is readily detected at the new SPB inner plaque but not at the outer plaque indicates that its asymmetry is not due to fluorophore maturation. Further, our data clearly demonstrate that SPB insertion is not the trigger for completion of outer plaque assembly.

As *γ*TC is recruited to the SPB by Spc72, we also examined Spc72 incorporation into the SPB during duplication using Spc42 to mark collectively the satellite stage and duplicated side-by-side SPBs (Figure 2 C). Like *γ*TC, we observed two localization patterns. Some cells showed single Spc72 label at the old SPB (Figure 2 C, a, arrowhead) while the majority of cells showed Spc72 label extending from the old SPB to the bridge, with label favoring one or the other location (Figure 2 C, b-d, big and small arrowheads, respectively) and a small proportion appearing to carry label at the bridge exclusively (Figure 2 C, e). This dual distribution was abrogated in *kar1Δ15* cells in which label remained at the old SPB (Figure 2 C, f). An identical result was obtained when Spc72-Venus localization was determined in reference to Nud1-mT2 label (Figure 2 D), which also displays two signals from the satellite stage and throughout duplication (Burns et al., 2015 and Supplementary Figure 3 C). Taken together, these data demonstrated the incomplete assembly of the new SPB outer plaque prior to SPB separation as depicted in the new model proposed in Figure 2 E.

Localization of Spc72 and Tub4 at and away from the bridge during the side-by-side stage was also assessed relative to the bridge component Kar1 (Supplementary Figure 3 D-H). By contrast to the dual distribution detected by linescan analysis for both Tub4 and Spc72 relative to Spc42 used as reference (Supplementary Figure 3 D-E), Kar1 was exclusively positioned between unseparated Spc42 signals (Supplementary Figure 3 F-G and also see Burns et al., 2015; Seybold et al., 2015). Three color imaging experiments using Venus-Kar1 in addition to Spc42-mCherry and Tub4-mT2 or Spc72-mT2 confirmed that the central peak of Spc72 or Tub4 co-localized with Kar1 at the bridge in unseparated SPBs (Supplementary Figure 3 H). Importantly, these experiments clearly showed that a significant fraction of both Spc72 and Tub4 are maintained at the old SPB, even in the presence of bridge-localized pools of each protein.

To confirm that the asymmetric localization of Spc72 and γTC arises during SPB duplication, we examined Spc72-Venus and Spc98-mT2 (another component of the *γ*TC) in *KAR1* and *kar1Δ15* cell populations released from a metaphase block imposed by repression of *CDC20*. This allowed us to follow synchronous populations proceeding unperturbed along the subsequent cell cycle to ensure we captured and staged all modes of localization during early landmarks of the spindle pathway (Figure 3). SPB duplication took place in cells with incipient buds (Figure 3 A, a, b, f), which began SPB separation 15 min later (Figure 3 B - D). SIM analysis of these synchronized cells confirmed the inherent asymmetry built into the SPB duplication cycle, the overall absence of Spc72 and Spc98 at the new SPB outer plaque, and a modest contribution to Spc72 label at the new SPB attributed to the bridge, as inferred from the nearly absolute asymmetry at onset of SPB separation observed in *kar1Δ15* cells. These findings validated our analysis using asynchronous cell populations (Figures 1 - 2). We additionally quantified Spc72 intensity at the old versus new SPBs upon separation in *kar1Δ15* cells to determine when Spc72 began to be incorporated to the new SPB outer plaque. As indicated by a significant drop in Spc72 asymmetry index at the transition from the ∼ 1-µm long stage, Spc72 began to accumulate in short spindles prior to anaphase onset (Figure 3 E).

**Figure 3.**
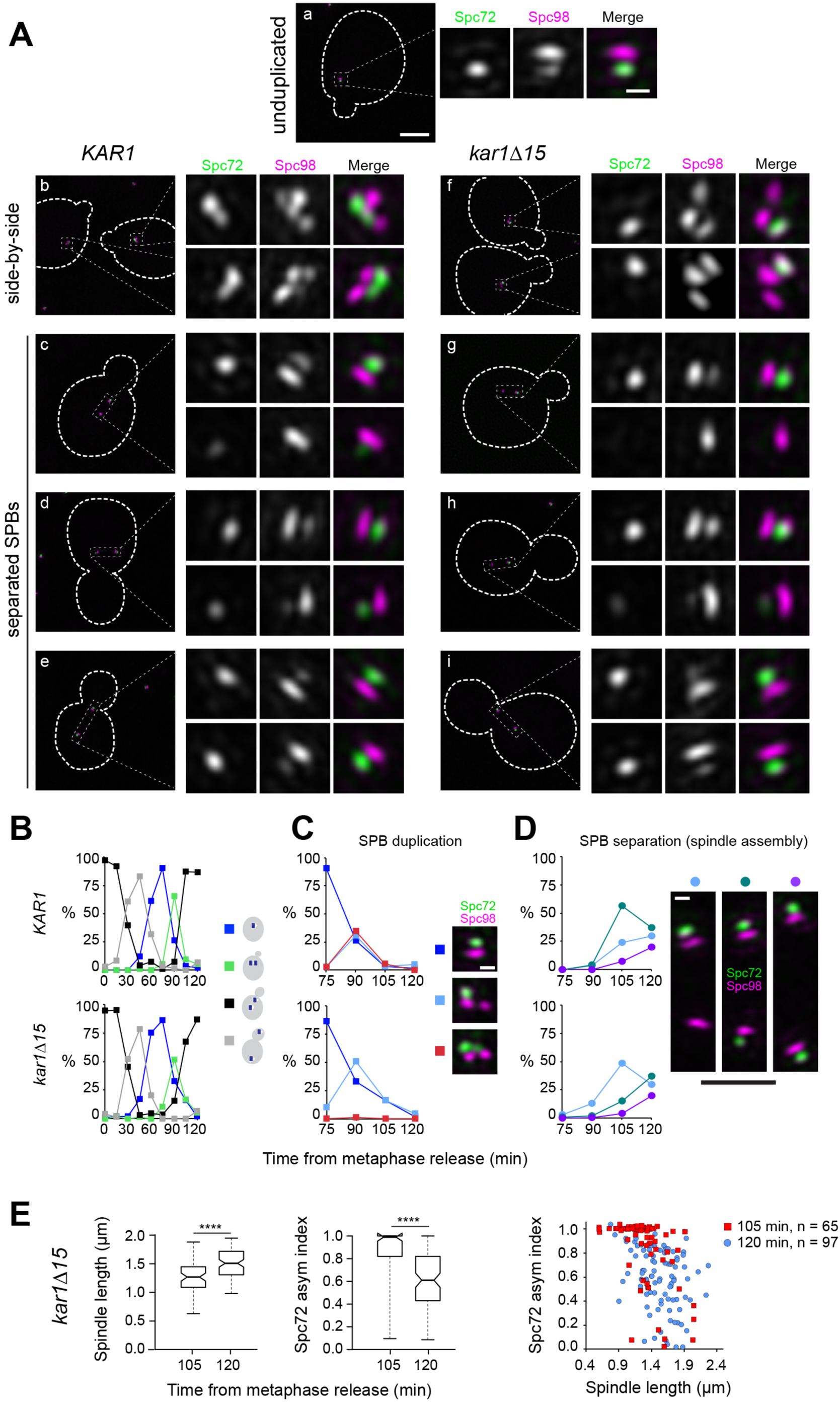
Spc72 accumulation at the new SPB outer plaque begins at the ∼ 1 µm-long spindle stage. (A-E) *KAR1* or *kar1Δ15* cells co-expressing Spc72-Venus (green) and Spc98-mT2 (magenta) were synchronized in metaphase using *P_MET3_-CDC20*, then released into the cell cycle. (A) Representative SIM images showing the distribution of Spc72-Venus (green) and Spc98-mT2 (magenta). Merged images with cell outlines (scale bar, 2 µm) and cropped images of SPBs by channel or merged (scale bar, 200 nm) are shown: (a) unduplicated, (b, f) side-by-side (c-e and g-i), separated SPBs. (B) Cell cycle progression in terms of the spindle pathway. (C-D) The localization of Spc72-Venus and Spc98-mT2 was quantitated at the indicated times in cells with side-by-side SPBs (C; scale bar, 200 nm) or separated SPBs (D; scale bars: white, 200 nm and black, 1 µm) Total number of cells analyzed by time point was 70, 85, 107, 123, 70, 114, 150, 198 and 125 (*KAR1*); 96, 160,103, 70, 128, 139, 320, 205 and 224 (*kar1Δ15*). (E) Boxplots for distribution of spindle length or Spc72 asymmetry index (left) and scattered plot for Spc72 asymmetry index as a function of spindle length at the indicated time points were generated by 3-D SIM analysis. ****, p < 0.0001 according to Mann Whitney two-tailed test.

In conclusion, throughout SPB duplication and the ensuing side-by-side stage, Spc72 and the cytoplasmic pool of *γ*TC are present at the old SPB outer plaque and the bridge but are absent from the new SPB outer plaque. This asymmetry persists at onset of SPB separation with the new SPB acquiring Spc72, and *γ*TC, as cells transit past the ∼1 µm-long spindle stage.

### S-phase CDK is required for Spc72 asymmetric recruitment

Our data show that Spc72 (and *γ*TC) bias to the old SPB at onset of spindle assembly stems from inherent asymmetry built into the SPB duplication cycle. The observation that tethering the *γ*TC-binding domain of Spc72 to Cnm67 (a component incorporated to the new SPB from the satellite stage) abrogates the delay in aMT acquisition by the new SPB during spindle assembly (Juanes et al., 2013) is consistent with this model. Intrinsic SPB asymmetry is also abolished in *cdc28-4 clb5Δ* mutant cells, with both SPBs exhibiting aMTs at onset of SPB separation (Segal et al., 2000). In order to determine whether this was also due to loss of Spc72 asymmetry, asynchronous populations of wild type, *clb5Δ*, *cdc28-4* and *cdc28-4 clb5Δ* mutant cells expressing Spc72-GFP were subject to quantitative imaging analysis. Using Spc42-CFP as a reference, the level of Spc72 at each SPB in cells carrying short spindles was normalized and an asymmetry index calculated (see Materials and Methods). Short spindles of wild type and *cdc28-4* mutant cells exhibited marked Spc72 asymmetry while a modest decrease was observed in *clb5Δ* cells (Figure 4 A). By contrast, asymmetry was significantly impaired in *cdc28-4 clb5Δ* cells (Figure 4 A-B), in agreement with the observation that this mutant nucleates aMTs at both SPBs at onset of spindle assembly (Segal et al., 2000).

**Figure 4.**
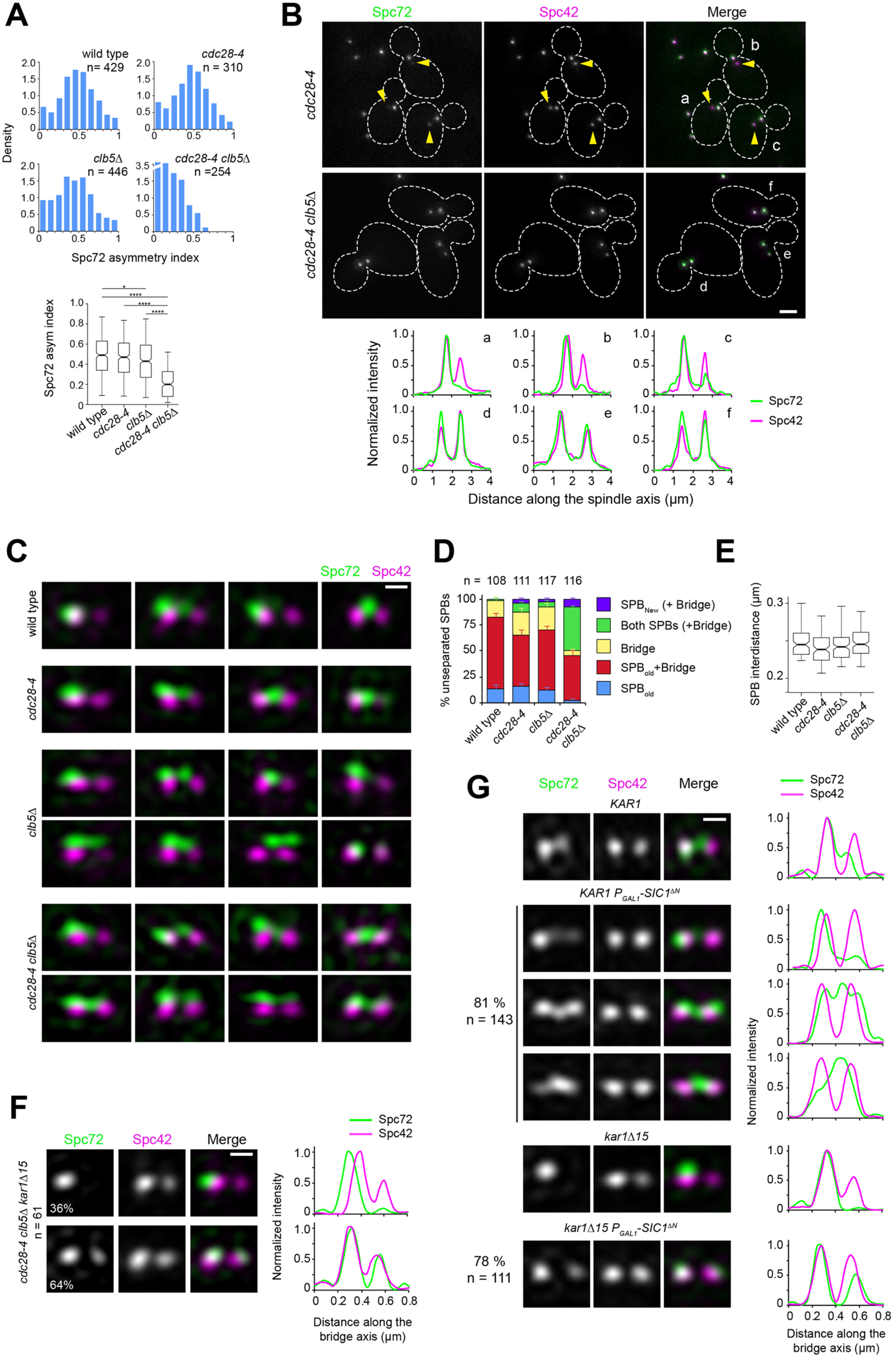
Spc72 asymmetric recruitment requires S-phase CDK. (A-B) The distribution of Spc72-GFP (green) and Spc42-CFP (magenta) was compared in wild-type, *clb5Δ*, *cdc28-4* and *cdc28-4 clb5Δ* cells grown at 23°C. (A) Histograms show the distribution of Spc72 asymmetry index in cells with short spindles (< 2.5 µm length) for each sample. Boxplots below depict the 5th, 25th, 50th, 75th and 95th centiles. Notches represent 95% CI of the median. * p = 0.0189 and **** p < 0.0001, according to Kruskal-Wallis and Dunn’s multiple comparison tests. (B) Representative wide-field fluorescence images and corresponding linescan analysis along the spindle axis showing Spc72 asymmetry in *cdc28-4*, but not *cdc28-4 clb5Δ* cells. Yellow arrowheads point to new SPBs weakly labelled by Spc72-GFP, underscoring asymmetry. Scale bar, 2 µm. (C-D) SIM analysis of unseparated SPBs of the indicated strains. (C) Representative 3D-realigned images of unseparated SPBs labeled by Spc72-Venus (green) and Spc42-mT2 (magenta). Loss of asymmetry occurred in *cdc28-4 clb5Δ* cells in which Spc72 recruitment at the new SPB outer plaque began prior to SPB separation. Scale bar, 200 nm. (D) Quantitation of modes of Spc72 localization in unseparated SPBs of the indicated strains. Error bars, standard error of the proportion. (E) Boxplot showing the distance between Spc42 foci in all cells tallied. 5th, 25th, 50th, 75th and 95th centiles are shown. Notches represent 95% CI of the median. (F) Representative images for the two modes of Spc72 localization observed in unseparated SPBs of *cdc28-4 clb5Δ kar1Δ15* asynchronous cells. Images were subject to realignment based on the Spc42-mT2 label. The corresponding linescans for fluorescence intensity along the bridge axis (3-px width; internally normalized) are also shown. (G) *KAR1* or *kar1Δ15 P_GAL1_-SIC1^ΔN^* cells (or control cells carrying a vector instead of pLD1) containing Spc72-Venus (green) and Spc42-mT2 were grown in synthetic-raffinose medium to early log phase at 25^°^C and induced at the same temperature by addition of 3% galactose. After 1.5-hour induction, SIM images were acquired and the SPBs in small-budded cells analyzed. Uniform cell cycle arrest was observed after 3.5 h. Representative SIM images of Spc72-Venus (green) and Spc42-mT2 (magenta) are shown with corresponding linescans (3-px width; internally normalized). Scale bar, 200 nm.

We further examined unseparated SPBs in these strains by SIM, as Clb5-Cdc28 is active during this stage of the SPB/cell cycle. A strong synergy was observed between the *cdc28-4* and *clb5Δ* mutations — in the double mutant, Spc72 was present at the new SPB already before SPB separation in more than 50% of cells analyzed (Figure 4 C-E). Additionally, introducing the *kar1Δ15* allele did not affect excess symmetry caused by *cdc28-4 clb5Δ* mutations confirming the involvement of the new SPB outer plaque (Figure 4 F). The genetic interaction was specific for *clb5Δ* and not shared by a *clb3Δ clb4Δ* mutant even though Clb3 and Clb4 are needed for timely SPB separation (Juanes et al., 2011; Segal et al., 1998 and Supplementary Figure 4). The requirement for S-phase CDK in restricting Spc72 to the old SPB during the side-by-side stage was further confirmed by SIM analysis of *KAR1* or *kar1Δ15* cells upon induction of an non-degradable version of the CDK inhibitor Sic1, which prevents entry into S phase and arrests cells prior to SPB separation (Figure 4 G). Taken together, these data suggest that S-phase CDK imparts correct spindle polarity by enforcing Spc72 asymmetry.

### CDK enforces Spc72 asymmetric recruitment via Nud1

To identify SPB components that might represent direct CDK targets involved in Spc72 asymmetric recruitment, we undertook a candidate-based approach guided by an SPB phospho-proteomic dataset (Keck et al., 2011) to generate a series of strains in which single SPB components (Cnm67, Kar1, Spc72 and Nud1) were replaced for mutant versions in which CDK consensus sites (minimal <u>S/T</u>-P or <u>S/T</u>-P-X-K/R) were cancelled by substituting the critical S or T for A (see Materials and Methods). The resulting strains were evaluated for Spc72 asymmetry in cells with short spindles by wide-field fluorescence microscopy. From those candidates, only cells expressing Nud1^7A^ in which positions S^21^, S^294^, T^388^, T^392^, S^469^, T^806^ and T^843^ were substituted by A to cancel seven CDK consensus sites, prompted a significant loss of Spc72 asymmetry *in vivo* (Figure 5 A-B). Importantly, the substitutions did not affect Nud1 expression level (Supplementary Figure 5 A). Mutation of these sites abolished phosphorylation by Clb5-Cdc28 *in vitro* (Supplementary Figure 5 B), with five of the seven sites also confirmed in subsequent global phosphoproteome studies of the SPB (Fong et al., 2018; Rock et al., 2013). Accordingly, Nud1^7A^ exhibited a reduction in bulk phosphorylation *in vivo* relative to wild type Nud1, based on analysis of mobility shift by western blot (Supplementary Figure 5 A and C). Consistent with the importance of intrinsic asymmetry in the establishment of spindle polarity and asymmetric fate, expression of Nud1^7A^ resulted in a modest increase in the reversed pattern of inheritance by which the new SPB entered the bud (29% versus 4% in cells expressing wild type Nud1, Supplementary Figure 5 D).

**Figure 5.**
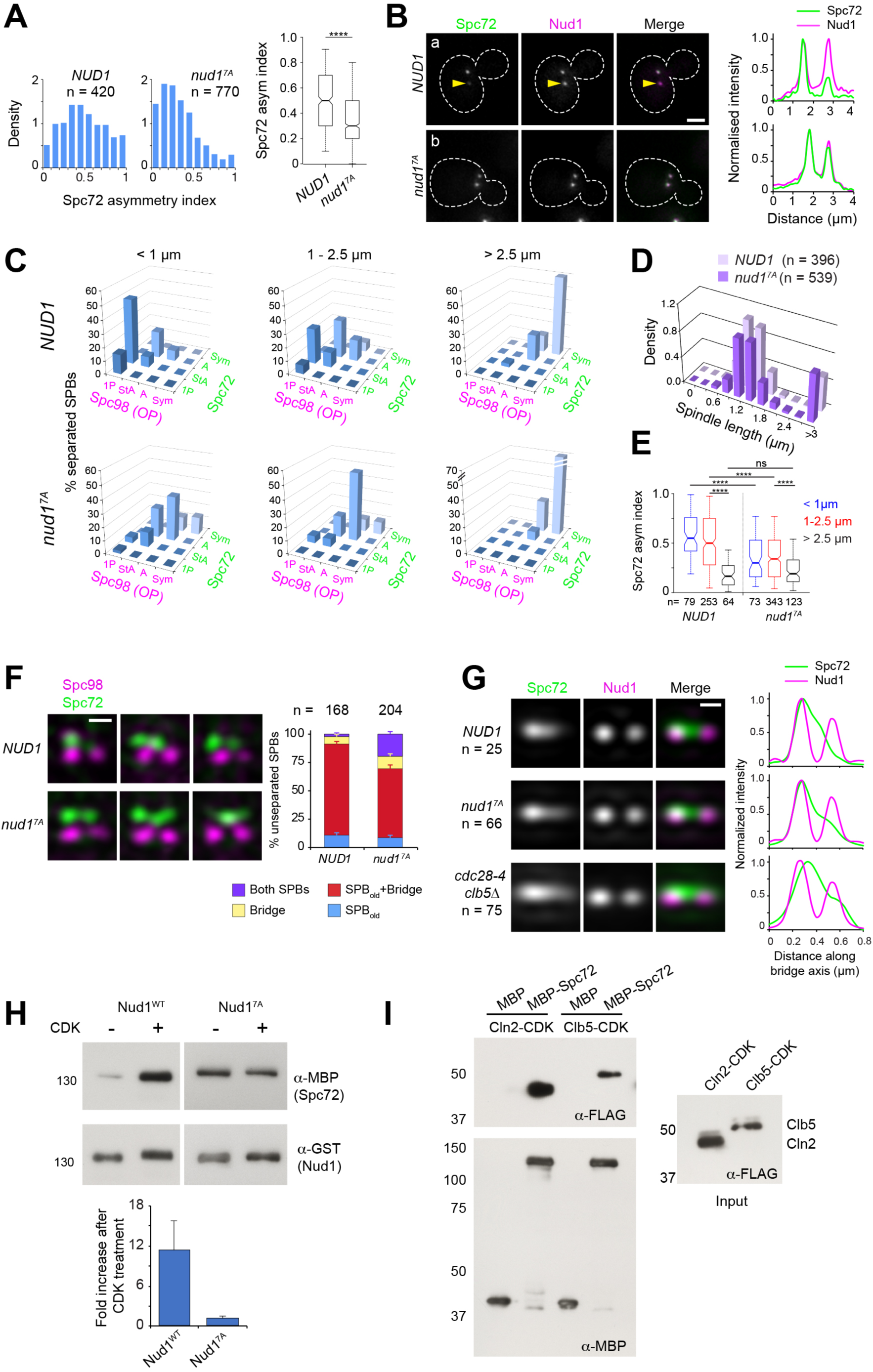
Substitutions cancelling CDK phosphorylation in Nud1 result in loss of Spc72 asymmetry. (A-B) Cells expressing wild type Nud1-CFP or Nud1^7A^-CFP along with Spc72-GFP were analyzed by wide-field fluorescence microscopy. (A) Histogram showing the distribution of Spc72 asymmetry index in each sample. **** p < 0.0001 according to the Mann Whitney two-tailed test. (B) Representative fluorescence images and corresponding linescan analysis along the spindle axis. Yellow arrowheads highlight asymmetric localization pointing to weaker label at the new SPB. Scale bar, 2 µm. (C) Distribution of Spc98-mT2 and Spc72-Venus by spindle stage was quantitated as described in Figure 1. (D) The spindle length distribution within the cell populations is shown. (E) Boxplot analysis showing Spc72 asymmetry index by spindle stage. 5th, 25th, 50th, 75th and 95th centiles are shown. Notches represent 95% CI of the median. ****, p < 0.0001 according to Kruskal-Wallis and Dunn’s multiple comparison tests. (F) (Left) Representative SIM images from cells with side-by-side SPBs to compare the distribution of Spc72 in *NUD1* versus *nud1^7A^* cells. In *nud1^7A^* cells, Spc72-Venus (green) was incorporated into the SPB outer plaque in a fraction of cells with side-by-side SPBs. Bar, 200 nm. (Right) Quantification of modes of Spc72 recruitment in unseparated SPBs of *NUD1* versus *nud1^7A^* cells. Error bars, standard error of the proportion. (G) Comparison of the extent of disruption of Spc72 asymmetry in *nud1^7A^* versus *cdc28-4 clb5Δ* cells assessed in averaged images of unseparated SPBs from asynchronous cell populations, compiled after 3D- realignment in reference to Nud1 label. The corresponding linescans for fluorescence intensity along the bridge axis (3-px width; internally normalized) are shown. (H) Effect of CDK phosphorylation of Nud1 vs Nud1^7A^ on Spc72 binding *in vitro*. (Top) A mixture of Cln2 and Clb5-CDK was used to phosphorylate GST-Nud1 or GST-Nud1^7A^ bound to beads. After extensive washing to terminate the kinase reaction, beads were incubated with identical amounts of purified MBP-Spc72 for 2 h followed by repeated washes. Eluted fractions were resolved in SDS-PAGE and the presence of Nud1 and Spc72 determined by western blot analysis. (Bottom) Plot representing average fold increase in Nud1-Spc72 binding upon CDK treatment from three independent experiments. Error bar, SD. (I) Ability of MBP-Spc72 to pull down Cln2 or Clb5 as part of CDK complexes *in vitro*. An equivalent amount of input (left, 5s exposure) was analyzed under identical conditions to the pull-down (right, 20s exposure).

If CDK-dependent phosphorylation acts through Nud1 to promote SPB functional asymmetry, then we would predict that *nud1^7A^* cells would display Spc72 symmetry at early stages of the spindle pathway, along the lines observed for a *cdc28-4 clb5Δ* mutant. To explore this possibility, *nud1Δ* strains complemented by *NUD1* or *nud1^7A^* and coexpressing Spc72-Venus and Spc98-mT2 were analysed by SIM. *nud1^7A^* cells already exhibited a marked reduction in Spc72 and Spc98 asymmetry at onset of SPB separation (Figure 5 C-E). Furthermore, we observed Spc72 recruitment to the new outer plaque in side-by-side SPBs of *nud1^7A^* cells (Figure 5 F). While reminiscent of that observed in *cdc28-4 clb5Δ* cells, the phenotype appeared less penetrant in *nud1^7A^* cells (Figure 5 F-G), suggesting that while Nud1 may be a key target, other CDK substrates may be involved.

To investigate the basis for CDK control of Spc72 asymmetry via Nud1, the ability of Spc72 to bind purified Nud1 or Nud1^7A^ after phosphorylation by yeast CDK was determined *in vitro*. CDK treatment resulted in a nearly 10-fold increase in pull-down efficiency for wild type Nud1 but exerted no effect on the extent of Spc72 binding to Nud1^7A^ (Figure 5 H). The interaction and its enhancement by CDK phosphorylation were retained by a Nud1 domain extending between amino acid positions 250-600 (Supplementary Figure 5 E-F). We hypothesized that this change in affinity might promote structural asymmetry if CDK activity inherently favored the old SPB outer plaque. To explore a mechanistic basis for such possibility, the interaction between G_1_/S cyclins and Spc72 was tested. Both Cln2 and Clb5 as part of CDK complexes were efficiently pulled down by purified Spc72 *in vitro* (Figure 5 I). It follows that changes in affinity induced by CDK phosphorylation of Nud1 may enforce Spc72 retention at the old SPB within a temporal window of opportunity initiated by Spc72 itself, thus reinforcing its bias to the old SPB.

### Orderly SPB assembly and correct γTC inner to outer plaque ratio require CDK

Spc72 recruitment at the new SPB outer plaque was preempted in greater than 50 % of unseparated SPBs of asynchronous *cdc28-4 clb5Δ* cells, a pronounced advancement compared to the recruitment of Spc72 to the new SPB outer plaque part way along spindle assembly observed in wild type cells. The *nud1^7A^* mutation also advanced Spc72 recruitment in unseparated SPBs, yet to a lesser extent (Figure 5), suggesting that additional CDK targets might be implicated. Yet, in view of the marked penetrance in *cdc28-4 clb5Δ* cells, it was of great interest to quantify the impact of CDK inactivation on the establishment of nucleation sites by analyzing Tub4-Venus localization in reference to Spc42-mT2 in *cdc28-4 clb5Δ* cells as done before for wild type cells (Figure 2). Surprisingly, this analysis pointed to comparable intensities of Tub4-Venus at inner versus outer plaque in this mutant (Supplementary Figure 6 A-B). We therefore proceeded to study Tub4-Venus localization relative to Nud1-mT2 instead, to assign inner and outer plaques at all stages without relying solely on the premise that the inner plaque corresponded to the more intense of the two Tub4 foci in a given SPB.

To determine the earliest point of *γ*TC recruitment at the new SPB outer plaque in *cdc28-4 clb5Δ* cells, the localization of Tub4-Venus was assessed in 3D-realigned SIM images of unseparated SPBs according to the Nud1-mT2 foci. Prior to the side-by-side stage, (paired Nud1-mT2 foci with a single Tub4 inner plaque focus), analysis of individual images indicated that Tub4-Venus distributed between the old SPB outer plaque (Figure 6 A, a-b; yellow arrowheads) and the bridge in wild type cells while present exclusively at the old SPB outer plaque in *kar1Δ15* cells (Figure 6 A, c-d, yellow arrowheads), in agreement with data presented in Figure 2. By contrast, 5 out of 12 *cdc28-4 clb5Δ* cells displayed a new Tub4 signal that colocalised with Nud1 at the presumptive new outer plaque before Tub4 presence at the new inner plaque (Figure 6 A, e versus f-g; hollow versus solid white arrowhead). A similar proportion of *cdc28-4 clb5Δ kar1Δ15* cells showed Tub4 at the new outer plaque in the absence of an inner plaque label (Figure 6 A, h versus i-j). These data suggested that *cdc28-4 clb5Δ* cells might preempt complete assembly of the new SPB outer plaque before the new inner plaque is fully established. Realigned images of cells after reaching the side-by-side stage (Tub4 at both inner plaques) were averaged for quantitation (Figure 6 B). Wild type *KAR1* or *kar1Δ15* cells subject to this protocol confirmed the intrinsically asymmetric side-by-side structure as documented in Figure 2 and Supplementary Figure 3. Yet, *cdc28-4 clb5Δ* (*KAR1* or *kar1Δ15*) mutant cells exhibited both significant accumulation of Tub4 at the new SPB outer plaque and an overall reduction in Tub4 IP:OP ratio at both SPBs. By analysing individual images used for these averages, the advanced Tub4 incorporation at the outer plaque relative to the inner plaque at the side-by-side stage was further apparent (Supplementary Figure 6 C). This advanced Tub4 addition at the outer plaque before inner plaque complete assembly followed Spc72 recruitment (Supplementary Figure 5 D).

**Figure 6.**
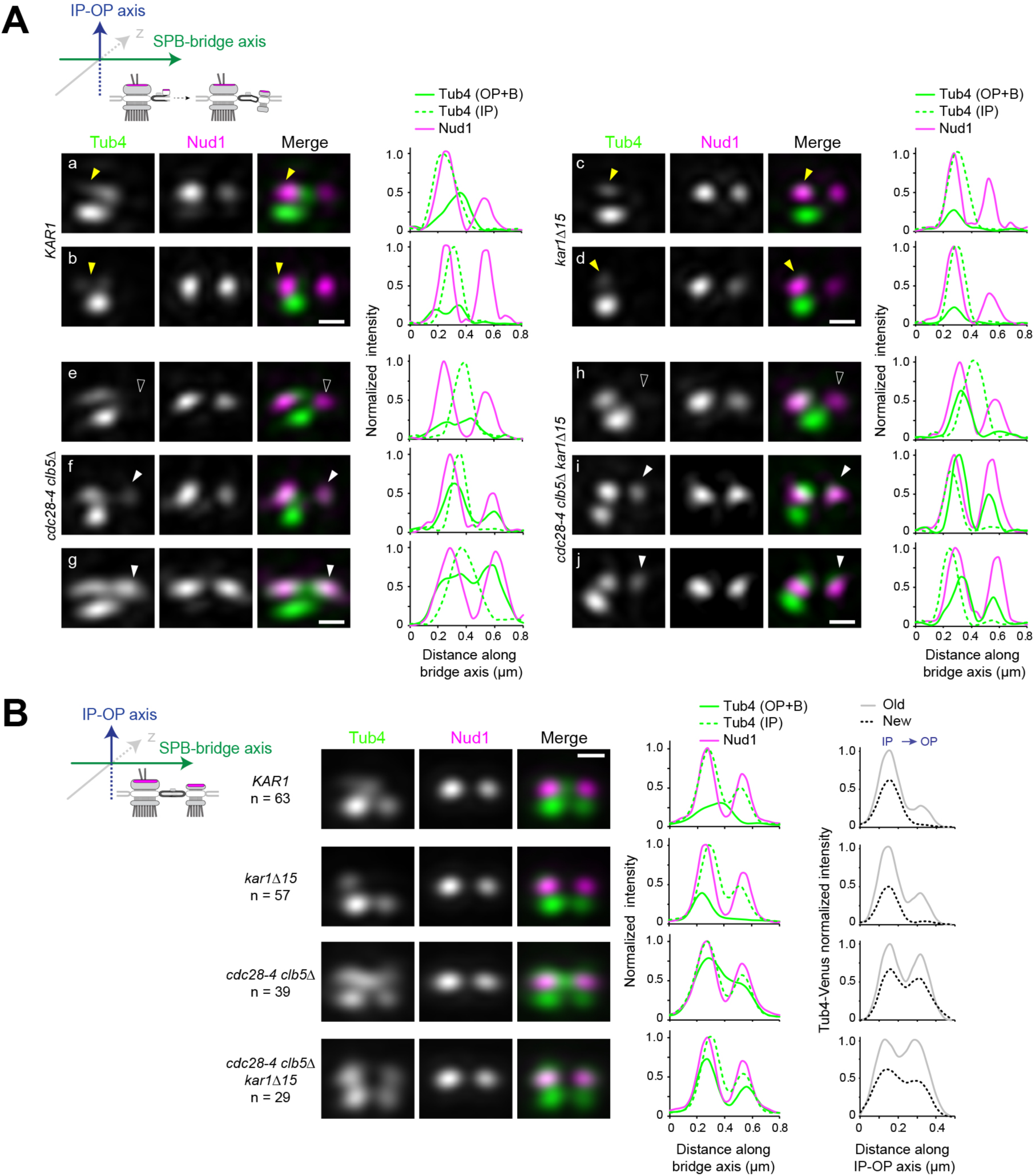
Tub4 IP:OP ratio and the sequence for new SPB assembly are perturbed in *cdc28-4 clb5Δ* cells. (A-B) SIM analysis of the indicated strains expressing Tub4-Venus and Nud1-mT2 as a reference for the outer plaque. Cartoons indicate the stage in each case and label used for 3D-realignment. Scale bar, 200 nm. (A) Individual SIM images and linescan analysis (1-px width; internally normalized) for the indicated strains corresponding to unseparated SPBs prior to nucleation establishment at the new SPB inner plaque, realigned with respect to Nud1-mT2. Tub4 colocalized partly with Nud1 at the old SPB in wild type cells (a-b; yellow arrowhead) or exclusively in *kar1Δ15* cells (c-d; yellow arrowhead). By contrast, in *cdc28-4 clb5Δ* and *cdc28-4 clb5Δ kar1Δ15* cells at this stage, Tub4 additionally colocalized with the new Nud1 focus (compare e with f-g or h with i-j; hollow versus solid white arrowheads). (B) Averaged images compiled after 3D-realignment with respect to Nud1-mT2 of side-by-side SPBs. Linescans for fluorescence intensity along the bridge and IP-OP axis are also shown (1-px width; internally normalized).

Contrary to what was observed in *cdc28-4 clb5Δ* cells, Tub4-Venus inner to outer plaque ratio in *nud1^7A^* cells was essentially unaffected (Supplementary Figure 6 E). Although the *nud1^7A^* mutation perturbed Tub4 asymmetry at the side-by-side stage in line with the advanced recruitment of Spc72 at the new SPB outer plaque (Figure 5), it was insufficient to reverse the normal sequence for addition of Tub4 recruitment during assembly of the new SPB (Supplementary Figure 6 F).

Taken together, inactivation of S-phase CDK in *cdc28-4 clb5Δ* cells not only abrogated asymmetry otherwise arising from the SPB duplication cycle but also reversed the orderly sequence of events required to build the new SPB, with concomitant impact on the balanced recruitment of the *γ*TC to each SPB face. These data demonstrated two separable contributions of CDK to the control of new SPB assembly impacting intrinsic asymmetry and correct SPB morphogenesis. First, CDK helps retain Spc72 at the old SPB outer plaque thus enforcing asymmetry in unseparated SPBs, at least in part, via phosphorylation of Nud1. Second, CDK addititionally imposes the sequential addition and the ensuing distribution of *γ*TC at the inner and outer plaques, most likely, by phosphorylation of other SPB components (see Discussion). These findings point to critical links between CDK temporal control, correct SPB morphogenesis and functional asymmetry.

## DISCUSSION

### SIM reveals that maturation and acquisition of nucleation capability are compartmentalized and proceed separately at inner and outer plaques of the new SPB

Historically, SPB assembly is thought to begin with the formation of the satellite, followed by addition of outer plaque components, including Nud1 and, perhaps by extrapolation, Spc72 and the *γ*TC (Adams and Kilmartin, 1999; Ruthnick and Schiebel, 2016; Winey and Bloom, 2012). Because mutants that disrupt SPB insertion into the nuclear envelope (i.e., *mps2-1*, *ndc1-1*) form a complete outer plaque that is competent for MT nucleation, SPB insertion is not required for outer plaque assembly (Winey et al., 1991; Winey et al., 1993). Yet, our current work challenges the idea that completion of the new outer plaque, specifically addition of Spc72 and *γ*TC, temporally precedes SPB insertion and even separation in wild type cells. Rather, these distal components are added later in the cell cycle as the new SPB matures part-way along spindle assembly.

EM analysis of SPBs and spindles and studies of fluorescently labeled α-tubulin show that the inner plaque nucleates 5-10-fold more MTs than the outer plaque (Winey and Bloom, 2012). Therefore, we anticipated to observe a correlative 5-10 IP:OP ratio in *γ*TC levels rather than the ∼2-fold difference we measured. Previous SIM was unable to estimate the overall proportion of SPBs presenting or lacking *γ*TC at the outer plaque, in part, because a fraction of SPBs were presumably aligned perpendicularly to the imaging plane, requiring resolution in the z axis greater than achievable (Lengefeld et al., 2018). Our use of a second label and three-dimensional realignment allowed us to include virtually all SPBs in our analysis, including unduplicated SPBs, side-by-side SPBs and SPBs in short and long spindles. Unlike direct stochastic optical resolution microscopy (dSTORM), which presumed a model for MT formation *in vivo* based on *in vitro* work, our analysis did not take into account anything other than fluorescence intensity, which increased during the cell cycle and when cell ploidy increased, suggesting it is a reliable readout of *γ*TC abundance. The discrepancy in *γ*TC levels compared to MT nucleation points to the fact that the availability of *γ*TC alone is not the sole regulator of MT formation in the cell. Further work will help elucidate the roles that the outer and inner plaque receptors, Spc72 and Spc110 respectively, the MT polymerase Stu2 (the budding yeast Dis1/TOG family member) and various post-translational modifications play in differential MT nucleation on each face of the SPB (Fong et al., 2018; Gunzelmann et al., 2018; Huisman et al., 2007; Keck et al., 2011; Lin et al., 2011; Lin et al., 2014). Our work illustrates an additional mode of regulation for aMTs, linked to SPB maturation that involves the addition of Spc72 and *γ*TC to the new SPB with a temporality emerging, at least in part, from CDK-dependent phosphorylation of Nud1.

### CDK contributions to the SPB cycle and spindle morphogenesis

CDKs promote multiple events along the spindle pathway (Avena et al., 2014; Byers and Goetsch, 1974; Chee and Haase, 2010; Elserafy et al., 2014; Fitch et al., 1992; Haase et al., 2001; Huisman et al., 2007; Jaspersen et al., 2004; Jones et al., 2018; Juanes et al., 2011; Liang et al., 2013; Lin et al., 2014; Ruthnick and Schiebel, 2016; Segal et al., 2000; Winey and Byers, 1993) beginning with the requirement for G_1_ cyclin (Cln)-Cdc28 in SPB duplication. Most *cdc28ts* alleles block cells at the satellite-bearing stage at the restrictive temperature. In turn, *cdc4ts* alleles (that prevent onset of Clb-dependent CDK activation) arrest cells after SPB duplication at the side-by-side stage (Byers and Goetsch, 1974). A number of SPB targets have been implicated but the precise molecular details are unknown (Jaspersen et al., 2004; Jones et al., 2018; Keck et al., 2011). CDK activity also ensures that SPB duplication occurs once and only once per cell cycle via Sfi1 — one of the best understood CDK molecular paradigms centered on an SPB component (Avena et al., 2014; Elserafy et al., 2014). Otherwise, most CDK phosphorylation sites characterised in SPB components have proven dispensable for viability (e.g. Huisman et al., 2007; Jaspersen et al., 2004; Lin et al., 2014, this study), with synergism observed, for example, when CDK sites are cancelled in multiple components (Jones et al., 2018). The extent of CDK-dependent phosphorylation at the SPB can be now appreciated by evaluation of phospho-proteomic datasets compiling phosphorylation sites at most SPB components (Fong et al., 2018; Keck et al., 2011; Lin et al., 2011; Rock et al., 2013) in the backdrop of phosphorylation by a number of additional cell cycle, and possibly other, protein kinases. However, it remains an ongoing challenge to link this complex regulatory landscape to function or any spatial restrictions tied into SPB age.

We previously implicated Clb5-Cdc28 in coordination of intrinsic SPB asymmetry with spindle morphogenesis on the basis of phenotypes observed in *cdc28-4 clb5Δ* mutant cells, a genetic setup designed to circumvent cyclin functional redundancy (Segal et al., 2000; Segal et al., 1998). Here our data indicates that Clb5-Cdc28 is required to impose correct temporality of Spc72 addition to the new SPB outer plaque. Indeed, *cdc28-4 clb5Δ* mutants exhibited a marked advancement of Spc72 recruitment in unseparated SPBs, a defect also apparent in cells overexpressing non-degradable Sic1, suggesting that while there might be functional overlap between Clns and Clb5, Cln-Cdc28 might be insufficient to sustain SPB intrinsic asymmetry. However, under unperturbed conditions we cannot exclude that both CDK complexes might contribute in this process, given the strong genetic interaction between *cdc28-4* (a G_1_ hypomorph *CDK* allele) and *clb5Δ* promoting this phenotype.

We found that substitutions cancelling CDK phosphorylation of Nud1 also advanced Spc72 recruitment to the new outer plaque prior to SPB separation. Nud1 plays dual roles as an SPB component. In addition to its structural role at the outer plaque in the recruitment of Spc72, Nud1 acts as a scaffold protein in the Mitotic Exit Network (MEN), a signalling pathway required for progression across the M/G_1_ phase boundary (Gruneberg et al., 2000; Rock et al., 2013). Previous studies suggested Nud1 as a putative CDK and Cdc14 target, the latter a phosphatase typically reversing CDK phosphorylation (Bloom et al., 2011; Park et al., 2008). Nud1 is also phosphorylated by Mps1, and, in connection with mitotic exit functions, by Cdc5 and Cdc15 (Keck et al., 2011; Maekawa et al., 2007; Park et al., 2008; Rock et al., 2013). Genetic analysis has previously implicated Nud1 in correct SPB inheritance but favored its MEN function in a mechanism linking the activation of the downstream MEN kinase Dbf2 at metaphase with polarized marking of the old SPB by Kar9 for bud-ward fate (Hotz et al., 2012). Yet, a more recent report (Campbell et al., 2019) demonstrated that Dbf2 activation is confined past anaphase onset to correctly regulate mitotic exit raising the possibility that the defective inheritance observed in a *nud1* mutant might reflect a structural contribution of Nud1 to the SPB demonstrated here. A previous study has also implicated both Nud1 and Spc72 in control of SPB inheritance during meiosis (Gordon et al., 2006).

Relative to a *cdc28-4 clb5Δ* mutant, *nud1^7A^* exhibited a less penetrant disruption of intrinsic SPB asymmetry. This suggests that CDK may exert its effect through further substrates that may include other components of the SPB, but also, more broadly via control of the activity of other protein kinases such as Mps1 and Cdc5, both dependent on CDK for their activity (Jaspersen et al., 2004; Mortensen et al., 2005; Rodriguez-Rodriguez et al., 2016). In addition, CDK also negatively regulates Cdc15 (Jaspersen and Morgan, 2000; Konig et al., 2010). Finally Clb5-Cdc28 also might antagonize Cdc14 by direct phosphorylation (Li et al., 2014), while the interplay between CDK thresholds and a broader constellation of phosphatases acting at the G_1_/S boundary is beginning to emerge (Arino et al., 2019; Martin et al., 2020).

Surprisingly, inactivation of S-phase CDK also affected the relative distribution of nucleation sites between the two faces of the SPB. Moreover, recruitment of *γ*TC was markedly advanced to the extent that Tub4 appeared at the new outer plaque before its incorporation to the new inner plaque in a fraction of duplicating SPBs, a reversal of the normal sequence of events in unperturbed cycling wild type cells. By contrast, *nud1^7A^* cells displayed excess symmetry in unseparated SPBs without otherwise decreasing Tub4 IP:OP ratio or altering the order in the assembly of the new SPB. These observations point to a critical contribution of CDK-enforced order through additional targets to govern correct *γ*TC partitioning, by promoting robust recruitment of *γ*TC first to the SPB inner plaque. Clb5-Cdc28 phosphorylation of Spc110 is critical for efficient *γ*TC recruitment in early S phase (Huisman et al., 2007; Lin et al., 2014). The *γ*TC assembles in the cytoplasm and nuclear import is mediated by a nuclear localization sequence in Spc98. Upon *γ*TC recruitment at the inner plaque (but not at the outer plaque), Spc98 undergoes Mps1 and CDK-dependent phosphorylation (Pereira et al., 1998), which is not essential for *γ*TC recruitment *per se* but might still promote retention and control nucleation activity and perhaps favor maturation of the new inner plaque ahead of the outer plaque. Tub4 also undergoes CDK and Cdc5-dependent phosphorylation that might be important for correct microtubule organization (Lin et al., 2011). In *cdc28-4 clb5Δ* cells, the combined effect of advanced Spc72 association with the new SPB and less efficient retention of *γ*TC by the inner plaque might cause roughly concurrent binding of *γ*TC to both faces of the SPB promoting an imbalance in the distribution of nucleation sites. By contrast, the *nud1^7A^* mutation does not affect *γ*TC regulation at the inner plaque, precluding this effect. In conclusion, CDK may enforce orderly SPB assembly that ensures both correct SPB morphogenesis and structural asymmetry.

### How does CDK promote Spc72 asymmetry via Nud1?

To understand the molecular link between Nud1 phosphorylation by CDK and Spc72 bias, the impact of CDK on the interaction between Nud1 and Spc72 was studied using purified components *in vitro*. Spc72 pull-down efficiency by phosphorylated wild type Nud1 as bait was enhanced approximately 10-fold over an unphosphorylated control. By contrast, CDK could not stimulate binding when Nud1^7A^ was used as bait. This change in affinity might translate into structural asymmetry if CDK activity were to inherently favor the old SPB outer plaque. In support of this possibility, we found that both Cln2 and Clb5 as part of CDK complexes interacted with purified Spc72 *in vitro*. These data suggest that CDK might preferentially associate with the old SPB, which contains Spc72 in G_1_, to impose structural and functional asymmetry later during the side-by-side stage and early spindle assembly. CDK phosphorylation of Nud1 once Clb5-Cdc28 becomes active, retains Spc72 at the old SPB, while Spc72 acquisition at the new SPB is inherently delayed. As Spc72 recruitment at the new SPB proceeds slowly due to the reduced affinity of Spc72 for non-phosphorylated Nud1, it might set in motion a positive feedback loop centered on Spc72 and CDK phosphorylation of Nud1 to accelarate completion of outer plaque assembly. Retention of Spc72 by the old SPB has been previously proposed although the mechanistic basis of how this might be achieved was not discussed (Lengefeld et al., 2018). Such a mechanism might effectively establish SPB identity on the basis of Spc72 inherited by the old SPB from the preceding cell cycle. In the absence of CDK phosphorylation of Nud1, however, Spc72 retention by the old SPB would be compromised during duplication and the side-by-side stage when it is partly redeployed to the bridge, and thus become readily shared by both SPBs. We therefore propose a built-in interplay between Spc72-driven recruitment of CDK at the old SPB and the ensuing enhancement of Spc72 binding affinity through Nud1 phosphorylation by CDK as a critical landmark event in the SPB duplication cycle linking SPB identity and age. The interplay between spatial control of distribution of the nucleation machinery and cell cycle-dependent phosphorylation may turn out a recurrent theme in centrosome control and beyond given the conservation of these nucleation systems and the protein kinase regulatory networks promoting maturation (Fu et al., 2015; Tovey and Conduit, 2018). Whether these mechanisms prove instrumental for cell cycle control of centrosome asymmetries associated with differential cell fate remains an exciting prospect.

## MATERIALS AND METHODS

### Yeast strains and genetic procedures

Yeast strains used in this study were derived from 15DaubA (Cepeda-Garcia et al., 2010; Juanes et al., 2011; Juanes et al., 2013; Richardson et al., 1989) or W303 (Burns et al., 2015) and are listed in Supplementary Table 1. Standard yeast genetic procedures and media were used throughout (Guthrie and Fink, 1991). Synthetic medium with supplements, or rich medium (YEP) contained 2% w/v dextrose, 3% w/v raffinose or galactose when indicated.

Strains carrying the *cdc28-4* allele in combination with deletions in cyclin genes have been previously described (Segal et al., 1998). Replacement of endogenous *KAR1* by the *kar1Δ15* allele (Pereira et al., 1999) was carried out by one-step disruption using pKS-kar1Δ15-URA3 or pKS-kar1Δ15-HIS2 (Juanes et al., 2013). Deletion of *NUD1* was carried out by one-step disruption using a targeted *KANMX* cassette produced by polymerase chain reaction (Juanes et al., 2013). Wild type *NUD1* gene or a mutated version in which seven CDK consensus sites were cancelled using nested PCRs (introducing S/T to A substitutions at positions 21, 294, 388, 392, 469, 806 and 843; referred to in the text as *nud1^7A^*) fused to CFP or HA_3_ were inserted as EcoRI-HindIII fragments into YIplac211 or YIp204, respectively. YIp211-NUD1-CFP or YIp211-NUD1^7A^-CFP were used for transformation of heterozygous *NUD1/nud1::KANMX* diploid cells. Linearization using StuI targeted integration at *URA3*. Following sporulation and tetrad dissection, haploid cells in which the *NUD1* deletion was rescued by the integrative constructs were selected. Haploid cells carrying YIp204-NUD1-HA_3_ or YIp204-NUD1^7A^-HA_3_ integrated at *TRP1* in the background of the *NUD1* deletion were similarly constructed. A similar strategy was implemented to construct strains expressing wild type or phospho-site mutant versions of Spc72-GFP, HA_3_-Kar1 or Cnm67-HA_3_ in the respective deletion backgrounds. Mutations encoded S/T to A substitutions at positions 232, 243, 552 and 566 of Spc72, positions 149, 216, 222 and 294 of Kar1 and positions 17, 72, 89, 103, 120, 121, 146 and 147 of Cnm67.

Endogenous tagging to produce C-terminal fusions to GFP, YFP, CFP, mCherry, or mTurquoise2 of Tub4, Spc72, Nud1, Spc42, Cnm67 and Spc98 in 15DaubA was carried out using tagging vectors previously described (Juanes et al., 2011; Juanes et al., 2013; Ten Hoopen et al., 2012). N-terminal tags fused to Kar1 and Spc110 were created by single-step replacement with recombinant cassettes targeted to the endogenous loci. W303-derived strains expressing fusions to fluorescent tags have been described in Burns et al. (2015). Strains transformed with pP*_MET3_*-*CDC20-LEU2* (a gift from Ethel Queralt) were used for synchronization by metaphase arrest and release as follows. Early log cell cultures grown in synthetic medium lacking methionine at 23°C were arrested by addition of 2.5 mM methionine in order to repress *CDC20* and incubation was continued for 3 h at 23°C. Uniform arrest was verified by microscopy. Cells were then harvested by centrifugation at 2000 rpm for 5 min at room temperature, rinsed twice and released by resuspending into synthetic medium lacking methionine at 23°C. Aliquots were collected every 15 min and processed for SIM. For inducible expression of non-degradable Sic1Δ^aa2-50^ under the *GAL1* promoter, cells were transformed with pLD1 (Noton and Diffley, 2000) linearized with ApaI. Mid-log cells grown in raffinose-containing synthetic medium at 23°C were induced by addition of 3% galactose and aliquots removed and processed every half hour. Uniform arrest (until cells exhibited single elongated buds) was achieved after 3.5 h.

### Wide-field fluorescence microscopy imaging and analysis

Images of cells coexpressing Spc72-GFP and Spc42-CFP (as reference) were acquired with a Nikon Eclipse E800 with a CFI Plan Apochromat 100x, NA 1.4 objective, a Chroma Technology CFP/YFP filter set and a Coolsnap-HQ CCD camera (Roper Scientific) (Guo and Segal, 2017) as five-plane Z-stacks of fluorescence images at a distance of 0.8 μm between planes with 2 x 2 binning, paired to a DIC image at the middle focal plane (Juanes et al., 2013). Stacks were processed into 2-color overlays of maximal intensity 2-D projections and analyzed with MetaMorph software (Molecular Devices). Linescans for fluorescence intensity along the spindle axis were generated with the line tool set to 3-pixel width and normalized intensity plots produced in Microsoft Excel. Integrated intensities were determined in a 7 x 7-pixel region and cell background subtracted for each channel. An asymmetry index was calculated as the absolute difference between the relative values at each SPB (i.e. Spc72/Spc42 intensity ratio) divided by the sum of the same values. The index ranged from 0 (absolute symmetry) to 1 (absolute asymmetry). Statistical analysis was carried out using GraphPad Prism and the open source package R. Boxplots depict 5th, 25th, 50th, 75th and 95th centiles with notches indicating the 95% confidence interval of the median.

### SIM imaging and analysis

Preparation of cell slides for SIM and image acquisition were according to Burns et al. (2015). Cells from early-log cultures grown at 22°C in synthetic complete medium supplemented with 0.1 g/L adenine were fixed in 4 % paraformaldehyde (Ted Pella) in 100 mM sucrose for 15 min at room temperature and washed twice in phosphate-buffer saline, pH 7.4. An aliquot of cells was resuspended in Dako mounting media (Agilent Technologies, #S3023), placed on a cleaned number 1.5 coverslip, covered with a cleaned glass slide then allowed to cure overnight at room temperature.

SIM images were acquired with an Applied Precision OMX BLAZE (GE Healthcare) equipped with an Olympus 60X 1.42 NA Plan Apo oil objective. Images were collected in sequential mode with two or three PCO Edge sCMOS cameras (Kelheim, Germany) for each acquisition channel. Color alignment from different cameras in the radial plane was performed using the color alignment slide from GE Healthcare. In the axial direction, color alignment was performed using 100 nm TetraSpeck beads (ThermoFisher, F8803). Reconstruction was accomplished with the softWoRx v6.52 software (GE Healthcare) according to manufacturer’s recommendations with a Wiener filter of 0.001. In most cases images are YFP/mT2 with 514 nm excitation for YFP and then 445 nm excitation for mT2. In some cases, we modified the protocol for mT2/YFP/mCherry acquisition with the mCherry acquired first and excited with the 568 nm laser. The dichroic in every case was 445/514/561 with emission filters at 460-485 nm, 530-552 nm and 590-628 nm for mT2, YFP and mCherry, respectively.

Three-dimensional analysis of SIM images was carried out using custom macros and plugins for the open source program FIJI (Schindelin et al., 2012). Plugins and source code are available for download at http://research.stowers.org/imagejplugins. First, spots corresponding to unseparated or separated SPB pairs were manually identified based on a reference label (typically Spc42 or Nud1) and sorted for analysis of the query label according to the experiment. If the maximum SPB intensity was in the first or last slice for either query or reference label, the SPB was excluded from further analysis. Next, for unseparated SPB pairs, the two spots in the reference channel representing the SPB were fitted to two 3D-Gaussian functions and realigned along the axis between these functions using [jay_sim_fitting_macro_multicolor_profile.ijm]. The higher intensity spot was assigned as the old SPB (Burns et al., 2015; Unruh et al., 2018). This realignment allowed the distribution of signal in the query channel to be analyzed against a single point of reference. For analysis of SPBs containing components of the *γ*TC as the reference channel, the inner and outer plaque signal were selected, with the stronger signal assigned to the inner plaque (see Figure S1). These images were fitted to two 3D Gaussian functions and subject to realignment. In some cases, after realignment, images were averaged and scaled as described previously (Burns et al., 2015), using [merge_all_stacks_jru_v1.ijm] then [stack_statistics_jru_v2.ijm]. Reconstructed images were scaled 4 x 4 with bilinear interpolation and presented as max projections in z over the relevant slices. After adjusting consistently brightness and contrast, and scaling to 8-bit image depth, cell images were incorporated into final figures using Adobe Illustrator.

SPB-SPB distances (referred to as SPB inter-distances) within cells were calculated after Gaussian fitting of the reference signal. Arbitrary boundaries along the spindle pathway were set according to the distribution of SPB inter-distances along the cell cycle measured in reference asynchronous cell populations labelled with Spc42-mTurquoise2 (Supplementary Figure 3) as follows: < 0.35µm, unseparated SPBs; <1µm-long spindles; 1-2.5 µm-long spindles; > 2.5 µm elongated spindles. Additionally, distance distributions were assessed for conformity in each experiment. In the special case of analyses of *γ*TC components with respect to the outer plaque component Spc72, SPBs were visually inspected and those in which inner and outer plaques overlapped vertically (as indicated by overlapping Spc72 and the sole *γ*TC signals) were excluded from analysis, due to limited resolution in the z axis to assess inner and outer plaque signals. As shown in Supplementary Figure 1E (using Nud1, a core outer plaque component as reference), only ∼12% of SPBs fell into this category. Linescan analysis along the IP-OP axis or the SPB-bridge axis was performed at 1-3- or 5-px width dependent on the experiment. When stated, normalization was carried out internally between maximal and minimal intensities in individual images or with respect to the indicated reference image in a set.

To directly compare Tub4 intensity in haploid and diploid cells, a haploid Tub4-mT2 Spc110-YFP strain was mixed with a Tub4-mT2/Tub4-mT2 diploid prior to imaging. After spot-fitting and realignment, images were assigned to the haploid or diploid group based on Spc110-YFP signal. Data from individual SPBs and averaged SPBs is shown.

### Protein production and purification

Recombinant MBP fusions to Nud1 or Nud1^7A^ were expressed from pMAL-c4X (New England Biolabs). Induction in Rosetta 2 cells (Novagen) was triggered with 0.5 mM IPTG followed by incubation at 16 °C overnight. Cells were then harvested by centrifugation 10 min at 5000 rpm at 4°C and washed with ice-cold H_2_O. Frozen cell pellets were stored at - 80°C. Bacterial cells were lysed by sonication and MBP fusions were purified onto amylose resin (New England Biolabs) as previously described (Ten Hoopen et al., 2012). To elute MBP-fusion proteins, the resin-bound fraction was resuspended in 500 µl of elution buffer (20 mM TRIS pH 7.5, 200 mM NaCl, 1 mM EDTA, 10 mM mercaptoethanol + 30 mM Maltose) and incubated on a roller for 5-15 min at 4°C. This was repeated and successive eluates tested for protein presence. Fractions were pooled together and subject to dialysis in HKEG buffer (20 mM HEPES, 50 mM KCl, 1mM EDTA, 5% Glycerol) using Maxi GeBaFlex-tube (Generon) at 4°C, concentrated by evaporation at 4°C and protein concentration evaluated by SDS-PAGE analysis against a BSA standard curve. Protein aliquots were stored frozen at −80°C.

Protein expression and purification from yeast was based on an auto-selection expression system previously developed (Geymonat et al., 2009). Cln2^ΔNt^ and Clb5^Δdb^, both stabilized versions of the cyclins Cln2 and Clb5, respectively (Hadwiger et al., 1989; Jacobson et al., 2000), were produced as GST or Twin-Strep-tag fusion proteins in the context of overexpressed Cdc28/Cks1. Briefly, PCR cassettes encoding the mutant cyclins along with gapped pMG1 or pMG6 expression vectors were used to transform the yeast expression host MGY70 (*MAT a ura3-1 trp1-28 leu2*Δ*0 lys2 his7 mob1::kanR pep4::LEU2 [URA3-MOB1])* or its derivative MGY139 (*MATα ura3 trp1 his3 leu2 lys2Δ0 pep4::LYS2 mob1::kanR cdc28::LEU2 [URA3-MOB1-CDC28])* by gap-repair. Auto-selection for the expression construct is achieved by passage of the resulting transformants onto FOA medium to select for loss of the resident *MOB1/CDC28*-carrying plasmid. Proteins were induced in YEP-galactose (1% final) for 8 hours at 30°C. Cells were harvested and proteins purified on glutathione beads according to Geymonat et al. (2009) or on Strep-Tactin XT resin (IBA) following manufacturer’s instructions. CDK protein complexes were eluted with 20 mM reduced glutathione or 50 mM biotin, respectively and dialyzed overnight at 4°C in buffer A (Tris pH 7.5 20 mM, NaCl 150 mM, DTT 0.5 mM, glycerol 10%). Co-purification of Cdc28 and Cks1 was confirmed by SDS-PAGE and cyclins quantified by Coomassie blue staining against a BSA standard curve. Constructs encoding GST-Nud1 or GST-Nud1^7A^ were also created by gap-repair of the pMG1 backbone in the yeast host MGY70. The resin-bound fraction was stored in 50% v/v glycerol at −20°C. MBP-Spc72 was expressed from the vector pMG3 in strain MGY853. MBP-Spc72 was eluted with 20 mM maltose and dialyzed o/n at 4°C in buffer A. Eluted proteins were quantified and stored at – 80°C.

### Western blot analysis, kinase and binding assays

Expression level of Nud1-HA_3_ and Nud1^7A^-HA_3_ was determined by western blot analysis of whole cell extracts as previously described (Ten Hoopen et al., 2012) using monoclonal antibody 12CA5 (Roche) at 1:1000 dilution and monoclonal antibody B-5-1-2 (Sigma) at 1:1000 dilution to detect *α*-tubulin as loading control.

4 µg of MBP-Nud1 or MBP-Nud^7A^ were incubated 30 min at 30°C in the presence of 40 µM ATP, 2 µCi [*γ*-^32^P] ATP (3000 mCi/mmol; 10mCi/ml, Perkin-Elmer) and 19 ng of purified Strep-tag-Clb5^Δdb^/Cdc28-as in kinase buffer (50 mM Tris pH 7.5, 10 mM MgCl_2_ and 1 mM DTT) in a final volume of 20 µl. When indicated, Cdc28-as was inhibited by addition of 5 µM 1NM-PP1. Reactions were stopped by addition of 5 µl of 5x Laemmli buffer followed by incubation at 95°C for 3 min. 10 µl- aliquots of phosphorylation reactions were analyzed on a 4-12% gradient SDS-PAGE gel and blotted to a PVDF membrane followed by autoradiography.

Equal amounts (∼3-5 µg) of GST-Nud1 and GST-Nud1^7A^ bound to beads (or Nud1^(250-600)^ and Nud1^4A(250-600)^) were incubated for 1 h at 37°C in a 200 µl reaction containing Strep-tag Clb5^Δdb^ and/or Cln2^ΔNt^/Cdc28/Cks1 (0.5-1 µg) and 1 mM ATP in kinase buffer (50 mM Tris pH 7.5, 10 mM MgCl_2_ and 1 mM DTT). Mock reactions were carried out in a 200 µl reaction containing kinase buffer only. Reactions were stopped by 3 washes of the beads with ice-cold washing buffer (50 mM Tris pH 7.5, 250 mM NaCl, 0.2% NP-40, 1 mM DTT). Beads were then resuspended in 300 µl of binding buffer (50 mM Tris pH7.5, 250 mM NaCl, 0.2% NP-40, 1 mM DTT, 1% BSA) containing 15 µg of purified MBP-Spc72 and incubated at 4°C for 2 h on a roller. Beads were washed 5 times with washing buffer and bound proteins were eluted with 20 mM reduced glutathione. Eluted proteins were analyzed on a 6% SDS-PAGE gel followed by a western blot using monoclonal anti-GST (Abcam) or anti-MBP (New England Biolabs) at 1:1000 dilution to detect GST-Nud1 and MBP-Spc72, respectively. Since MBP-Spc72 and GST-Nud1 have approximate molecular mass of 130 kDa, eluted material was loaded in duplicate gels for western blot analysis and probed with *α*-MBP or *α*-GST antibodies, respectively.

**Table 1.**
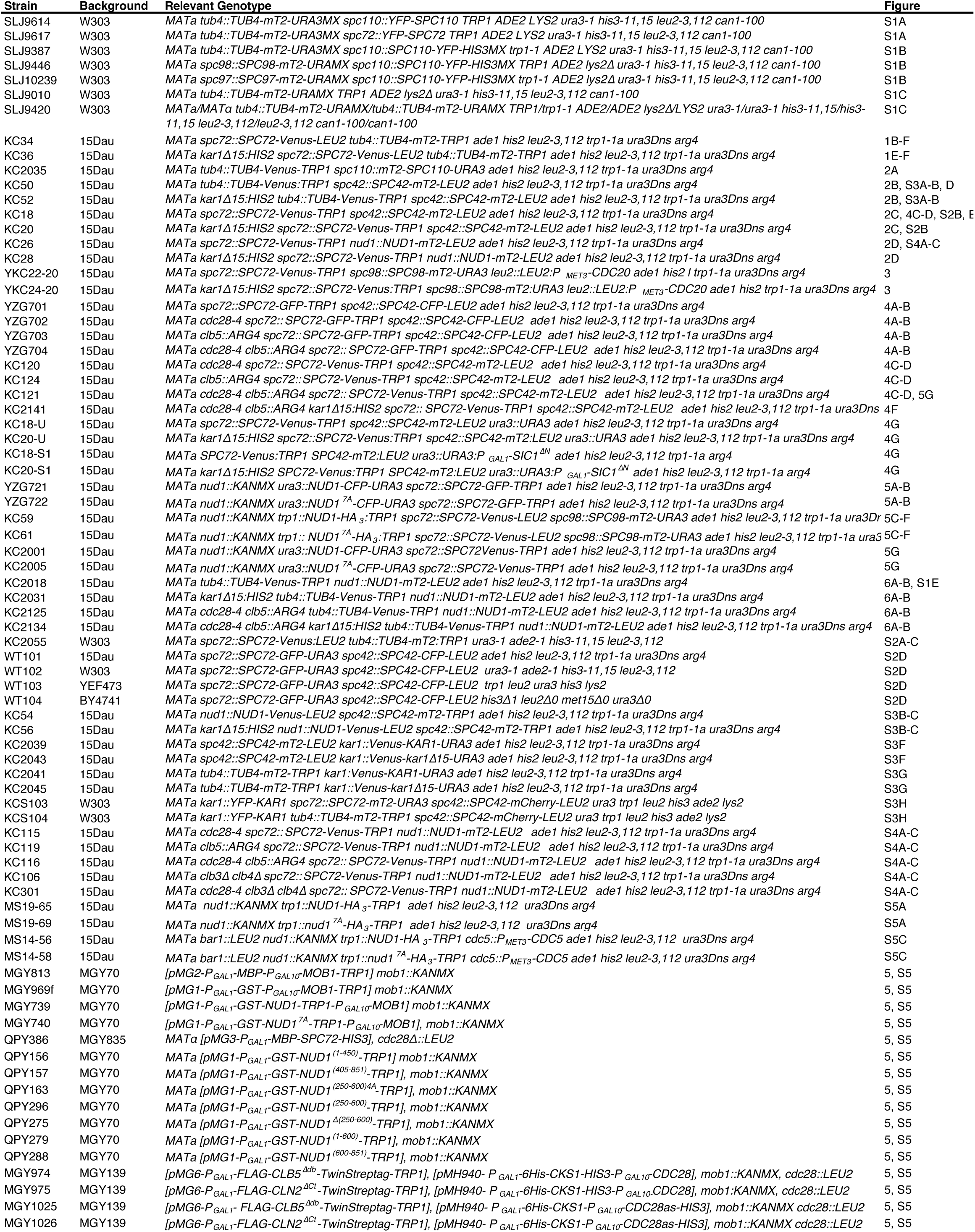
Yeast strains used in this study

## Acknowledgements

We are grateful to Mark Winey for sharing data prior to publication and to Mark Longtine John Kilmartin and Ethel Queralt for their gift of strains and constructs. We thank Jennifer Gardner, Shannon Burns, Luisa Capalbo, Alexandra Van Hall-Beauvais and Miriam Scarpa for contributing data to this project and Nikola Dzhindzhev for fruitful discussions. Research reported in this publication was supported, in part, by the Stowers Institute for Medical Research and the NIH-NIGMS under award number R01GM121443 (to SLJ), the Department of Genetics, University of Cambridge (to MS) and by CSC Cambridge International Scholarships (to QP and ZG). Original data underlying this manuscript can be downloaded from the Stowers Original Data Repository at http://www.stowers.org/research/publications/LIBPB-1526. The authors declare no competing financial interests.

## SUPPLEMENTARY FIGURE LEGENDS

**Figure S1.**
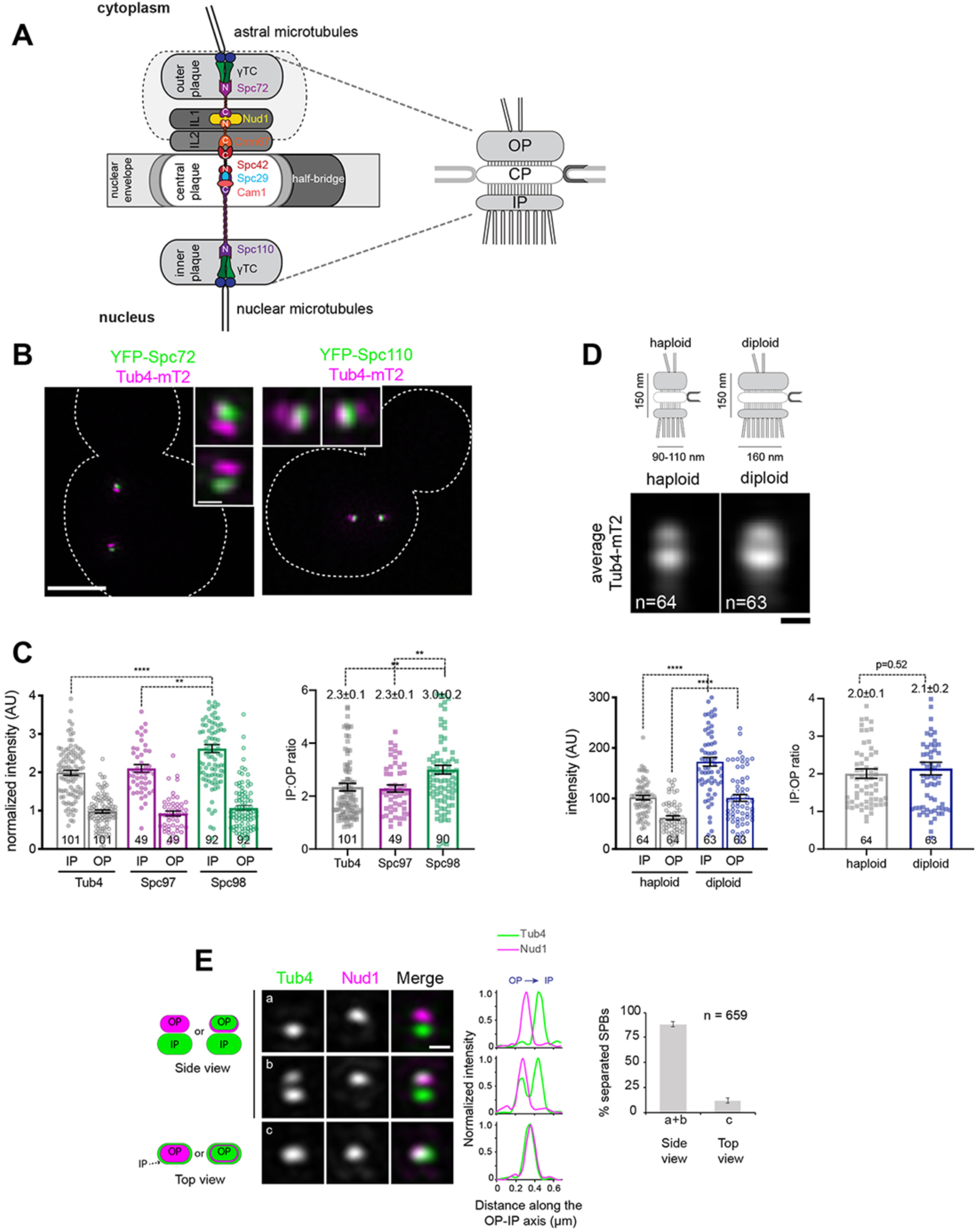
Quantitative detection of γTC by SIM shows SPB size scaling and roles for the outer plaque. (A) Cartoon outlining key layers and sublayers making up the SPB (left) and their correspondence to the summary structure (right) used to describe landmark events in the SPB duplication pathway (Figure 2) and SPB 3-D realignment throughout. (B) SIM images of cells containing Tub4-mT2 (magenta) and YFP-Spc72 or YFP-Spc110 (green), which mark the SPB outer and inner plaque, respectively. Insets show SPB region. Bar, 2 µm and 200 nm (inset). (C) SIM images of strains containing Spc110-YFP and the indicated γTC component fused to mT2 were analyzed by fitting spots corresponding to the inner and outer plaques. Each individual value is plotted along with the mean, standard error and number of SPBs analyzed (n). Ratios were derived from intensity measurements. (D) Schematic showing the inner, central and outer plaques of the SPB along with the half-bridge. The average width and height based on EM measurements of haploid and diploid cells is shown (Adams and Kilmartin, 1999; Byers and Goetsch, 1974). Haploid and diploid cells containing Tub4-mT2 were imaged by SIM and analyzed as in (C). In addition, an average image was also created. n, number of cells. ****, p<0.0001; **, p<0.01. Note that in both (C) and (D), there is heterogeneity within individual SPBs at both the inner and outer plaque. Importantly, SIM was able to detect quantitative differences in inner and outer plaque intensity that scaled proportionally with SPB size, indicating that it can be used to assay changes at the SPB. (E) SIM images of wild type cells expressing Tub4-Venus and Nud1-mT2 were used to score the overall proportion of individual SPBs presented in side view (a or b) and top view (c). Error bars, standard error of the proportion. Scale bar, 200 nm.

**Figure S2.**
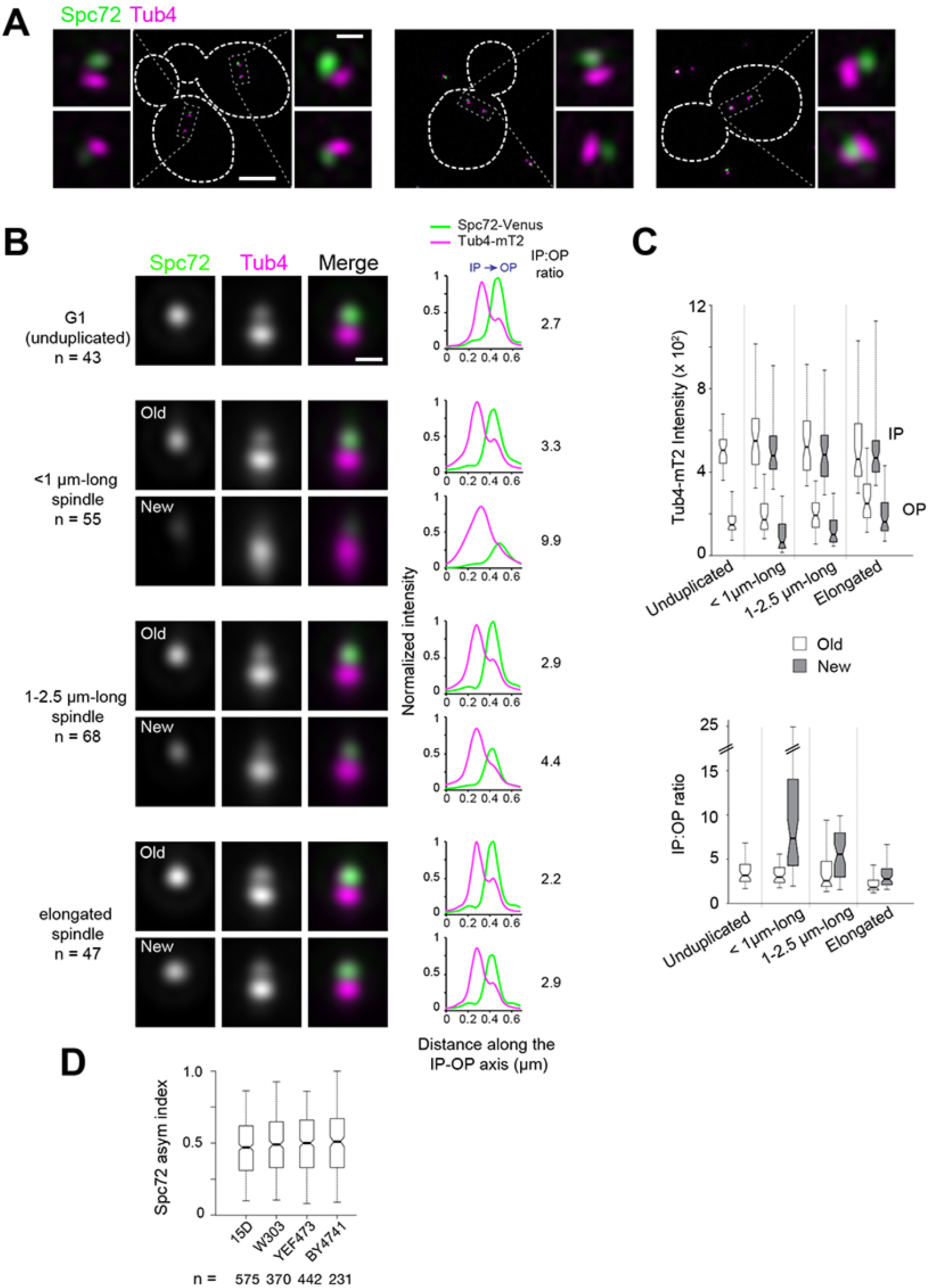
Spc72 and Tub4 asymmetric localization in the W303 yeast background. A W303-derived yeast strain containing Spc72-YFP (green) and Tub4-mT2 (magenta) was imaged by SIM. (A) Representative SIM images of cells with short spindles. Merged images with cell outlines (scale bar, 2 µm) and cropped merged images for each SPB (scale bar, 200 nm) are shown. (B-C) As in Figure 1, SIM images of individual SPBs were aligned using Tub4-mT2 as a reference and SPBs were assigned to a cell cycle and spindle stage using bud morphology and SPB inter-distances. (B) Averaged image from realigned SPBs in each class. Scale bar, 200 nm. The number of SPBs is indicated. Linescan analysis (5-px width; normalized relative to maximal intensity at the old SPB from elongated spindles) shows the intensity and localization of Spc72-YFP as well as the distribution of Tub4-mT2 at the IP and OP. The IP:OP ratio is based on the Tub4-mT2 signal. (C) In addition to averaging, the intensity of Tub4-mT2 at the inner and outer plaques was measured in individual images. Values are plotted by stage and SPB identity (top); IP:OP intensity ratios (bottom) for the same dataset are also shown. (D) Boxplot for the distribution of Spc72 asymmetry index in short spindles (<2.5 µm-long) for asynchronous cultures in the indicated wild type yeast backgrounds from wide-field images. Boxplots depict the 5th, 25th, 50th, 75th and 95th centiles. Notches represent 95% CI of the median.

**Figure S3.**
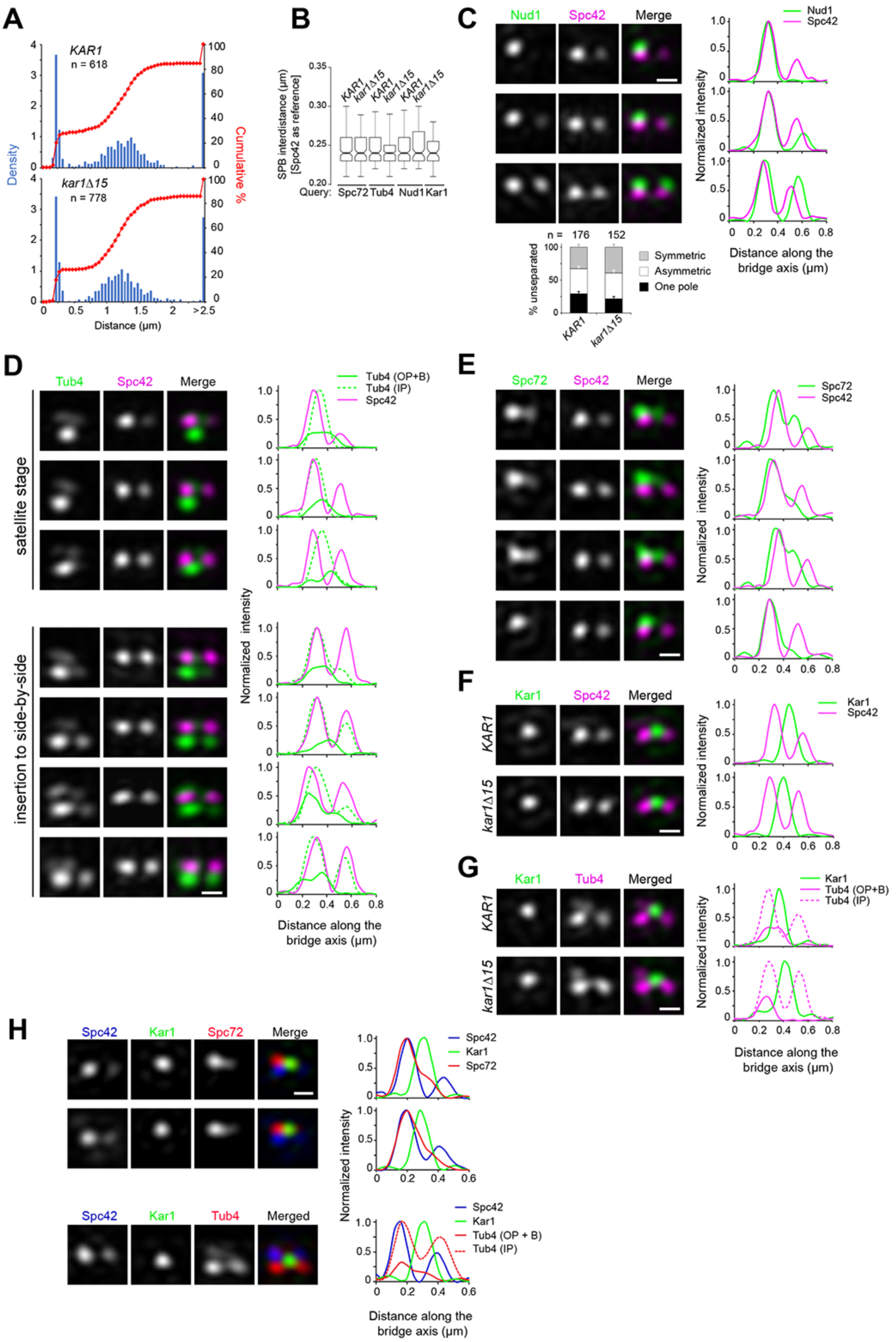
SIM analysis of intrinsic SPB asymmetry during SPB duplication. (A) Distribution of SPB inter-distances using Spc42-mT2 as reference in asynchronous *KAR1* or *kar1Δ15* cell populations analyzed by 3D-SIM. These values were used to classify SPBs into unseparated SPBs (< 350 nm), SPBs in short spindles (<2.5 µm) and SPBs in elongated spindles (>2.5 µm). (B) Boxplot for distribution of SPB inter-distances in all unseparated SPBs selected for analysis from asynchronous cell populations in Figures 2, 3 and Supplementary Figure 3 C. In all cases, analysis was performed after Spc42-mT2 labels (reference channel) were subject to 3D-Gaussian fit and realignment, as described in the Materials and Methods. (C) Nud1-Venus (green) and Spc42-mT2 (magenta) were localized in *KAR1* versus *kar1Δ15* cells by SIM. Three representative images show modes of label in unseparated SPBs. Consistent with the previously shown order of satellite assembly (Burns et al., 2015), Nud1 did not mark the presumptive satellite in approximately 25% of cells containing two Spc42 signals, but otherwise significantly accumulated at the new SPB toward the side-by-side stage. This is quantitated below. (D-H) A subset of cytoplasmic nucleation sites coincide with Kar1 localization in unseparated SPBs. (D-F) Comparative localization of Tub4-Venus, Spc72-Venus, Venus-Kar1 or Venus-Kar1Δ15 (in green) relative to Spc42-mT2 (magenta). Representative SIM image projections were obtained after 3D-Gaussian fit and realignment using Spc42-mT2 as reference. The corresponding linescan for fluorescence intensity along the bridge axis is also shown. When indicated, separate profiles were obtained along inner and outer plaques (1 pixel-width, parallel to the bridge axis). As quantified in Figure 2 B-C, Tub4-Venus or Spc72-Venus exhibited dual localization at the cytoplasmic face of duplicated SPBs by distributing between the old SPB outer plaque and the bridge. By contrast, Venus-Kar1 or Venus-kar1Δ15 was confined to the space between Spc42-mT2 foci, representing the exclusive localization to the bridge. (G) Partial overlap between Tub4-mT2 (magenta) and Venus-Kar1 (green) signals at the bridge was abolished when cells expressed instead Venus-Kar1Δ15. Scale bar, 200 nm. (H) Three color SIM of unseparated SPBs containing YFP-Kar1 (green), Spc42-mCherry (blue) and Spc72-mT2 or Tub4-mT2 (red). Linescan analysis was carried out along inner and outer plaques (1 pixel-width, parallel to the bridge axis). Scale bar, 200 nm.

**Figure S4.**
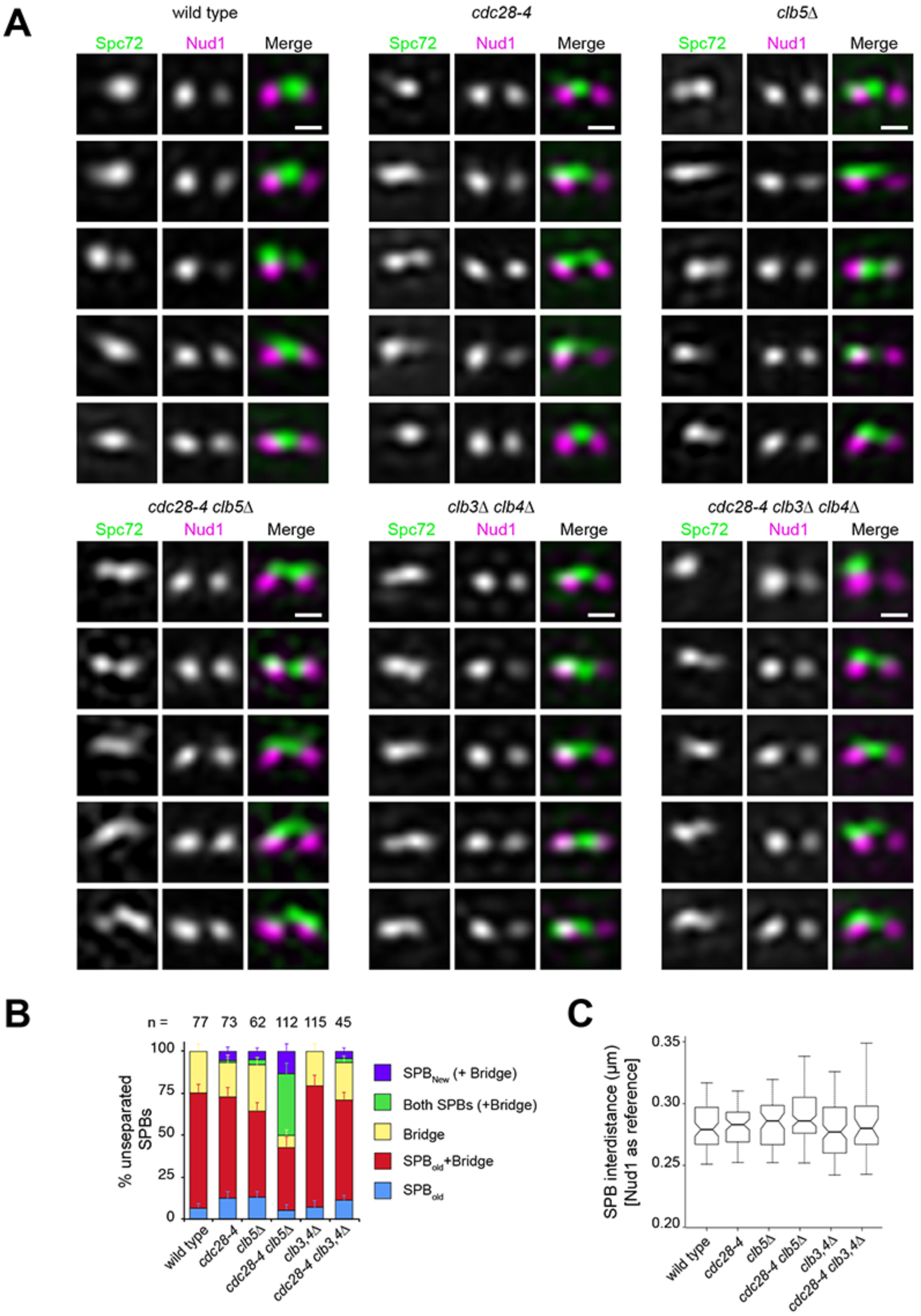
S phase CDK inactivation anticipates Spc72 recruitment at the new outer plaque in unseparated SPBs. Wild-type and *cdc28-4* and cyclin mutant strains containing Spc72-Venus (green) and Nud1-mT2 (magenta) were imaged by SIM and subject to 3-D Gaussian fit and realignment, in reference to Nud1, to determine the distribution of Spc72 relative to Nud1. (A) Representative SIM images showing unseparated SPBs. Scale bar, 200 nm. (B) Quantitation of Spc72 distribution in all strains analyzed. Bars, standard error of the proportion. (C) Boxplot for distance between Nud1 unseparated foci for all cells analyzed. Boxplots depict the 5th, 25th, 50th, 75th and 95th centiles. Notches represent 95% CI of the median.

**Figure S5.**
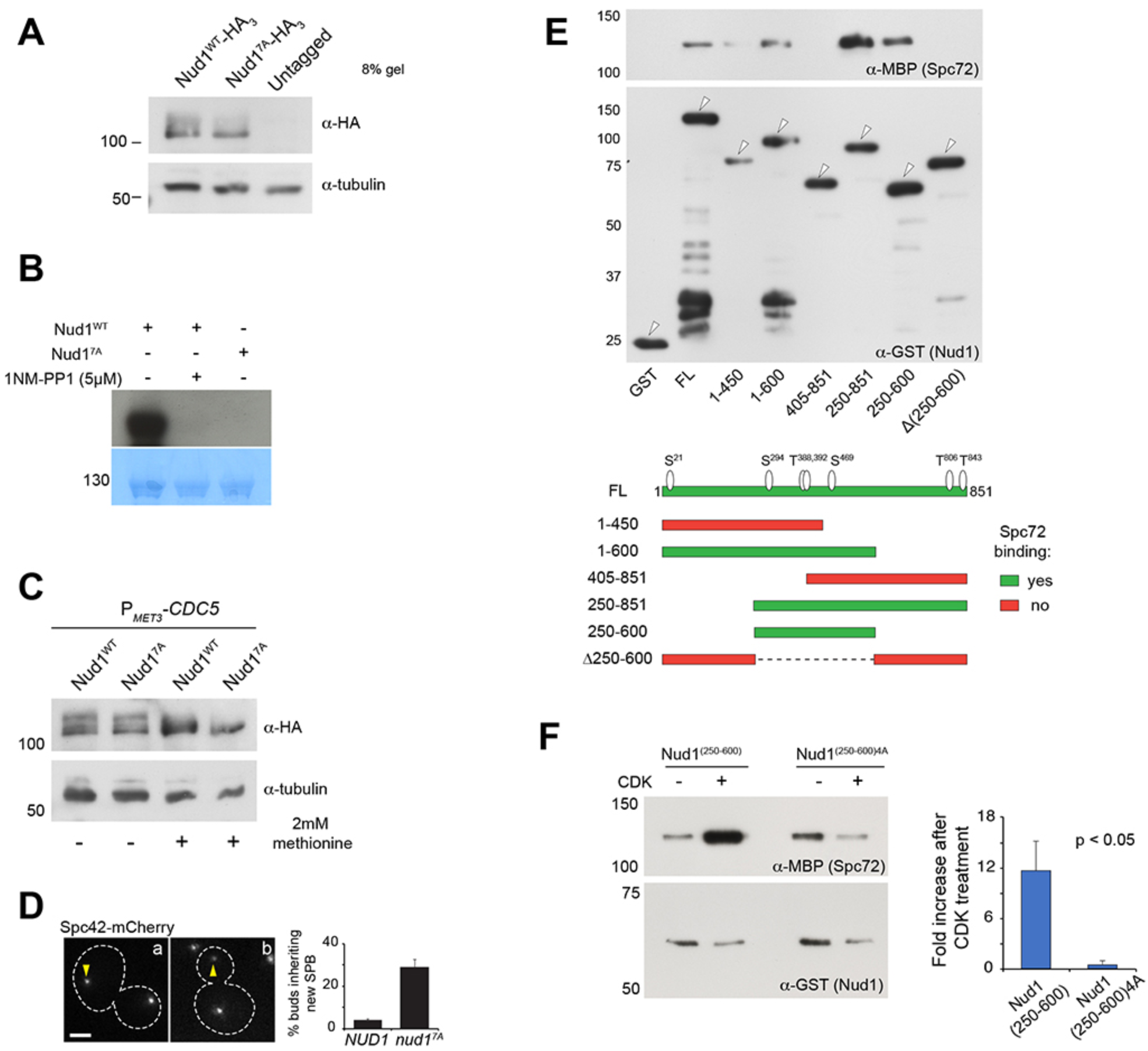
Nud1 and Nud1^7A^ characterization *in vivo* and *in vitro*. (A) Nud1^WT^-HA_3_ and Nud1^7A^-HA_3_ levels assessed by western blot analysis of cell extracts prepared from the parent strains analyzed in Figure 5. (B) Kinase assay using 4 µg of purified MBP-Nud1^WT^ or MBP-Nud1^7A^ and 19 ng of purified Twin-Strep-Clb5^Δdb^/Cdc28- as/Cks1. 5 µM 1NM-PP1 was added to inhibit Cdc28-as. Reactions were resolved by SDS-PAGE and blotted onto a PVDF membrane (bottom) that was exposed to film overnight (top) (C) CDK sites in Nud1 contribute to bulk phosphorylation *in vivo*. Western blot analysis of whole cell extracts from cells expressing Nud1^WT^-HA_3_ or Nud1^7A^-HA_3_ before or after depletion of Cdc5 under the control of the repressible *MET3* promoter. Cdc5 also contributes to Nud1 mobility shift by phosphorylation. Residual shift of Nud1 following Cdc5 repression was abolished upon substitutions cancelling CDK phosphorylation sites. (D) Proportion of *NUD1* versus *nud1^7A^* cells in which the new SPB was inherited by the bud. Mean of three counts of 200 cells is shown, error bars SD. (E) A Nud1 domain between amino acid positions 250-600 was necessary and sufficient for Spc72 binding *in vitro*. Previous yeast two-hybrid studies suggested that the Nud1 C-terminus (residues 405-852) associated with Spc72 (Gruneberg et al., 2000). (F) Spc72 pull down by Nud1(250-600) showing enhanced binding following CDK phosphorylation as described for full-length Nud1 in Figure 5 J.

**Figure S6.**
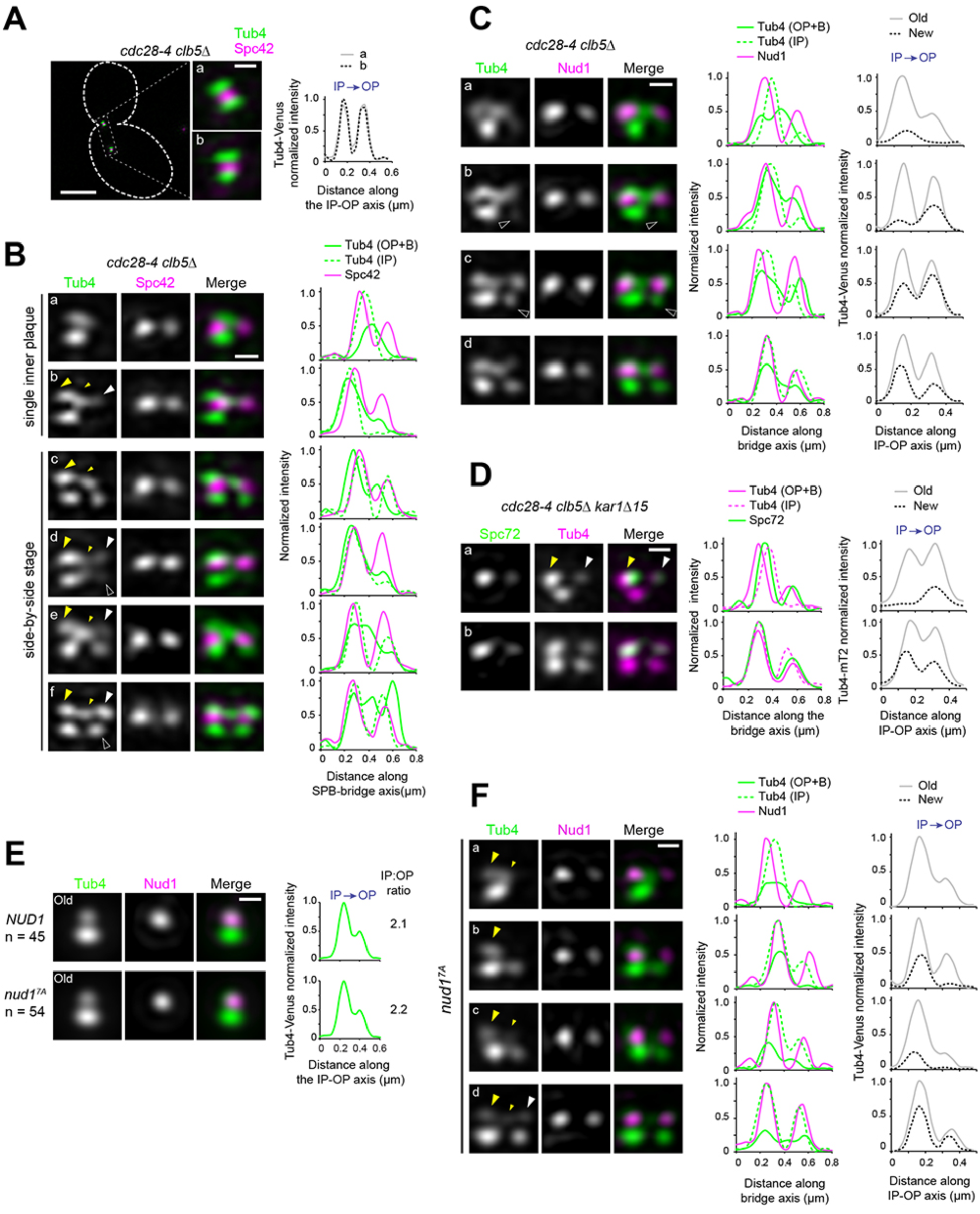
Nud1^7A^ anticipates Spc72 recruitment without otherwise affecting the Tub4 IP:OP ratio. (A-B) Representative SIM images for Tub4-Venus localization relative to Spc42-mT2 in *cdc28-4 clb5Δ* cells showing a marked decrease in the IP:OP ratio. (A) Merged image with cell outline (scale bar, 2 µm) and cropped merged images of SPBs (scale bar, 200 nm) corresponding to a cell with a short spindle. Linescan analysis (3-px width) along the IP-OP axis for each SPB is also shown. (B) Realigned SIM images of unseparated SPBs in reference to Spc42-mT2. Tub4 distributed between the old SPB outer plaque and the bridge during duplication (big and small yellow arrowheads, respectively) but was also recruited to the new SPB outer plaque. Early in duplication, some cells appeared to incorporate Tub4 at the new outer plaque before the inner plaque was complete (b, white arrowhead) but others preserved the correct order (c). Hollow white arrowheads point to weaker IP signal relative to OP at the new SPB, also suggestive of perturbed order. Linescan analysis along inner or outer plaque (1px-width) is also shown. (C) Selected realigned images of unseparated SPBs in *cdc28-4 clb5Δ* cells expressing Tub4-Venus and Nud1-mT2 from the dataset used to generate the average image shown in Figure 6 D alongside linescan analysis. Cells began to add Tub4 to the new inner plaque before (a) or after (b) incorporation of Tub4 to the outer plaque. Accordingly, cells at the side-by side-stage could exhibit a weaker Tub4 signal at the new inner plaque relative to the new outer plaque (c, hollow arrowhead). (D) Spc72 recruitment at the new SPB outer plaque may be detected in SIM images of *cdc28-4 clb5Δ kar1Δ15* expressing Spc72-Venus and Tub4-mT2. During duplication, Spc72-Venus was present at the old SPB outer plaque (yellow arrowhead) but additionally marked the presumptive new SPB outer plaque (white arrowhead) in 5 out of 15 cells showing a single inner plaque focus of Tub4-mT2 (a). Aberrant IP:OP ratio also trailed the premature incorporation of Spc72 at the side-by-side stage (b). (E) Tub4 IP:OP ratio in SPBs of *NUD1* versus *nud1^7A^* cells carrying short spindles (<2.5 µm-long). SIM averaged images of old SPBs obtained after 3D-Gaussian fitting and realignment using Tub4-Venus as reference. Scale bar, 200 nm. The number of SPBs is indicated. Linescan analysis for Tub4-Venus intensity along the IP-OP axis is shown. In addition to averaging, boxplots with the distribution of Tub4-Venus intensities at the inner and outer plaques measured in individual images and the corresponding IP:OP ratio are also presented. (F) Representative images of unseparated SPBs in asynchronous *nud1^7A^* cells following 3D-Gaussian fitting and realignment using Nud1^7A^-CFP as reference. Tub4-Venus distributed between the old SPB and bridge during duplication (a-c; big and small yellow arrowheads, respectively) but was prematurely added to the new SPB outer plaque at the side-by-side stage (after a Tub4 focus was established at the new inner plaque) without otherwise perturbing the IP:OP ratio (d).

